# A purine salvage bottleneck leads to bacterial adenine cross-feeding

**DOI:** 10.1101/2023.10.17.562681

**Authors:** Ying-Chih Chuang, Nicholas W. Haas, Robert Pepin, Megan Behringer, Yasuhiro Oda, Breah LaSarre, Caroline S. Harwood, James B. McKinlay

**Author notes:** Corresponding author: 1001 E 3^rd^ Street, Bloomington, IN 47405, USA. Department of Plant Pathology, Entomology, and Microbiology, Iowa State University, Ames, Iowa, USA.

## Abstract

Diverse ecosystems host microbial relationships that are stabilized by nutrient cross-feeding. Cross-feeding can involve metabolites that should hold value for the producer. Externalization of such communally valuable metabolites is often unexpected and difficult to predict. Previously, we fortuitously discovered purine externalization by *Rhodopseudomonas palustris* by its ability to rescue growth of an *Escherichia coli* purine auxotroph. Here we found that an *E. coli* purine auxotroph can stably coexist with *R. palustris* due to purine cross-feeding. We identified the cross-fed purine as adenine. Adenine was externalized by *R. palustris* under diverse growth conditions. Computational models suggested that adenine externalization occurs via passive diffusion across the cytoplasmic membrane. RNAseq analysis led us to hypothesize that accumulation and externalization of adenine stems from an adenine salvage bottleneck at the enzyme encoded by *apt*. Ectopic expression of *apt* eliminated adenine externalization, supporting our hypothesis. A comparison of 49 *R. palustris* strains suggested that purine externalization is relatively common, with 15 of the strains exhibiting the trait. Purine externalization was correlated with the genomic orientation of *apt* orientation, but *apt* orientation alone could not explain adenine externalization in some strains. Our results provide a mechanistic understanding of how a communally valuable metabolite can participate in cross-feeding. Our findings also highlight the challenge in identifying genetic signatures for metabolite externalization.

## Introduction

Cross-feeding between microbes is central to processes ranging from biogeochemical cycles to the human microbiome (1). Although widespread, much remains unknown about the molecular mechanisms underlying metabolite cross-feeding via externalization of metabolites into the extracellular space. Here we use externalization as a catch-all definition for any mode of metabolite externalization (2). One of the most perplexing aspects of cross-feeding is the externalization of metabolites that hold value not just for the recipient but also for the producer. Externalization of such communally valuable metabolites could pose a fitness disadvantage for the producer, especially if the trait is exploited by a non-reciprocating neighbor. Nonetheless, there are many examples of cross-feeding of communally valuable metabolites (1-5).

To study cross-feeding, many researchers use synthetic microbial communities, or cocultures. Cocultures allow the researcher to preserve ecological aspects of interest while maintaining control over environmental and genetic parameters (6). Enforcing obligate cross-feeding of essential nutrients can ensure coexistence and reproducible outcomes. However, one cannot control or predict all the ways that microbes will interact. Understanding how microbes interact within synthetic communities is important to correctly interpret results from these increasingly popular research systems, and to predict microbial interactions in nature.

We previously designed a coculture to be dependent on the exchange of essential carbon and nitrogen (7). Specifically, we paired fermentative *Escherichia coli* and phototrophic *Rhodopseudomonas palustris* in an anaerobic minimal medium with glucose and N_2_ gas as the sole carbon and nitrogen sources, respectively (Fig 1A, top). *E. coli* fermented glucose to organic acids that served as an essential carbon source for *R. palustris. R. palustris* reciprocated by excreting ammonium (NH_4_ ^+^), derived from N_2_, in a process called N_2_ fixation. NH_4_ ^+^ excretion relied upon engineered mutations that resulted in constitutive activity of the N_2_-fixing enzyme nitrogenase (strain Nx, NifA* mutation) (7).

**Fig 1.**
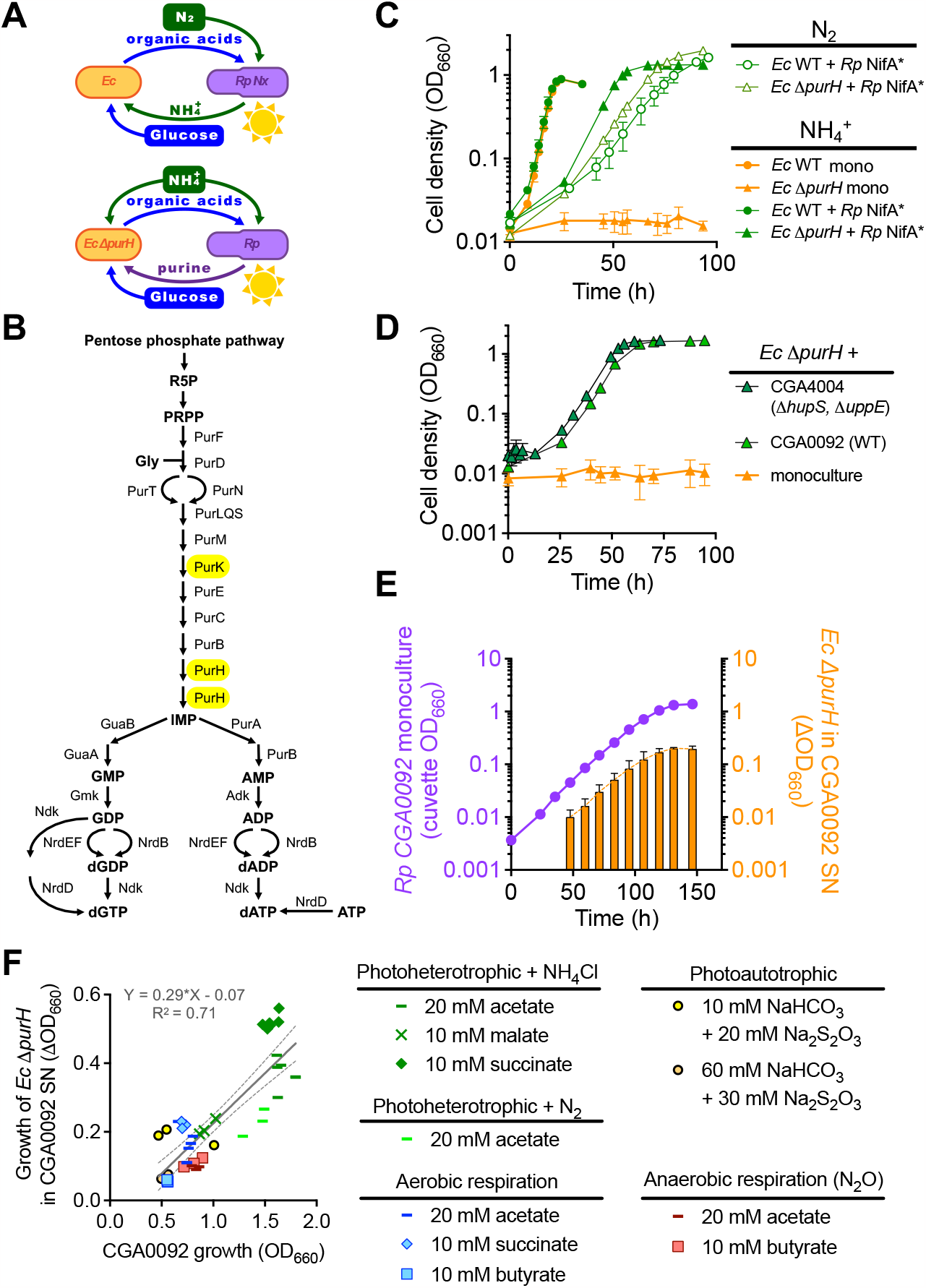
Wild-type *R. palustris* (*Rp*) CGA0092 supports *E. coli* (*Ec*) purine auxotroph growth across diverse growth conditions. **A.** Hypothesized critical cross-feeing interactions when N_2_ is the nitrogen source (top) or when *E. coli* is a purine auxotroph (bottom). **B**. *E. coli* de-novo purine synthesis pathway. **C**. Growth curves for *E. coli* monocultures and cocultures with *R. palustris* NifA^*^ with either N_2_ or NH_4_Cl as the sole nitrogen source. **D**. Growth curves for *E. coli* Δ*purH* in coculture with *R. palustris* strains having a wild-type *nifA* gene. NH_4_Cl was the sole nitrogen source. **E**. Growth of *E. coli* Δ*purH* in supernatants from *R. palustris* CGA0092 monocultures. **C-E**. Error bars, SD; n=3. Some error bars are smaller than the symbols. **F**. Growth of *E. coli* Δ*purH* in supernatants from stationary-phase CGA0092 monocultures grown under various growth conditions. Each data point represents a single biological replicate. Linear regression (gray line) +/- 95% confidence intervals (dashed lines) was applied to all samples across all conditions. SN, supernatant.

In working with the above coculture, we uncovered unanticipated layers of interaction. One notable example was revealed when we assessed the contribution of each *E. coli* gene to fitness in monoculture versus coculture using a transposon mutant library (8). *E. coli* purine synthesis genes were dispensable in coculture but not in monoculture, suggesting that *R. palustris* externalized purine(s) at quantities that can sustain a purine auxotroph. Here, we characterize the molecular basis for purine externalization by *R. palustris* and the potential for purine cross-feeding in both synthetic and natural communities.

## Materials and Methods

### Bacterial strains

Strains and 16S rRNA accession numbers are listed in Table S1. *R. palustris* CGA4004 has a Δ*hupS* mutation that prevents H_2_ oxidation and a Δ*uppE* mutation that prevents cell aggregation (9). *R. palustris* Nx (CGA4005) additionally carries a *nifA*^*^ mutation that results in NH_4_^+^ excretion under N_2_-fixing conditions (7). *E. coli* MG1655 (10) deletion mutants were made via lambda Red recombination (11) using deletion constructs amplified from *E. coli* KEIO mutants (12). Plasmids and primers are listed in Tables S2 and S3, respectively.

The adenosine phosphoribosyltransferase expression vector, pBBPgdh-apt, was generated using *E. coli*-mediated assembly (13) with *E. coli* NEB10β. Transformants were grown on lysogeny agar with 20 µg/ml gentamycin (Gm). Colony PCR was used to screen for correct plasmids, followed by verification by Sanger sequencing. pBBPgdh-apt and pBBPgdh were transformed into CGA0092 by electroporation (14) and selected on photosynthetic medium (PM) agar (15) with 10 mM disodium succinate and 100 μg/ml Gm.

### Growth conditions

Anaerobic media were prepared by bubbling with N_2_, then sealing with rubber stoppers and aluminum crimps prior to autoclaving. All anaerobic cultures were grown in 10-ml volumes in 27-ml anaerobic test tubes except for cocultures used to collect time-course data, which used 60-ml volumes in 150-ml serum vials.

*R. palustris* and *E. coli* were recovered from 25% glycerol frozen stocks on PM agar with 10 mM disodium succinate and lysogeny agar, respectively. Single colonies were used to inoculate starter cultures. *R. palustris* starter cultures were grown anaerobically in minimal M9-derived coculture medium (MDC) (7) with 20 mM acetate and 10 mM NH_4_Cl. *E. coli* starter cultures were grown aerobically in lysogeny broth (LB), with 30 μg/ml kanamycin (Km) when appropriate. To prepare *E. coli* for cocultures or bioassays, 0.2 ml of starter culture was centrifuged, and cell pellets were washed twice in 1 ml MDC. Cocultures were inoculated with 0.1 ml of *R. palustris* culture (diluted in MDC) and the washed *E. coli* cell suspension to an initial optical density (OD_660_) of ∼0.003 each. Cocultures were grown in MDC with 25 mM glucose, 10 mM NH_4_Cl, and 1% v/v cation solution (1 mM MgSO_4_ and 0.1 mM CaCl_2_), unless indicated otherwise. Photoautotrophic conditions used the indicated NaHCO_3_ and Na_2_S_2_O_3_ concentrations in place of organic carbon. Anaerobic chemotrophic conditions were supplemented with 0.1 mM NaNO_3_ and 100% N_2_O as described (15). All cultures were grown in horizontally oriented tubes or serum vials at 30°C with shaking at 225 rpm. Where indicated, light was provided by a 45 W halogen bulb (430 lumens).

### Analytical procedures

Cell densities were measured via turbidity (OD_660_) using a Genesys 20 spectrophotometer (Thermo-Fisher). Glucose and fermentation products were measured using high-performance liquid chromatography (HPLC; Shimadzu) as described (16).

### Invasion-from-rare assays

Cocultures of *E. coli* Δ*purH* and *R. palustris* CGA4004 were started from a range of initial frequencies for a total initial cell density of ∼ 10^6^ colony forming units (CFU) / ml. Samples were taken upon inoculation and after 5 days to determine frequencies by CFUs on aerobic lysogeny agar for *E. coli* and on anaerobic N_2_-fixing agar (PM without ammonium) with 10 mM succinate for *R. palustris*. Change in frequency was calculated as: (*E. coli* / (*E. coli* + *R. palustris*))_final_ – (*E. coli* / (*E. coli* + *R. palustris*))_initial_ (17).

### Metabolite extraction

Samples (1 ml) of *R. palustris* supernatant were spiked with internal standards of ^13^C5-adenosine (97%) and ^15^N3-dCMP (98%) (Cambridge Isotope Laboratories). Supernatant compounds were then extracted with four volumes of 1:1 v/v acetonitrile/methanol. The extraction mixture was incubated at -20°C for 20 min before centrifugation at 18,400 x *g* at 4°C for 15 min. Supernatants were transferred to 15-ml conical tubes, lyophilized, and stored at -20°C.

Intracellular compounds were extracted from *R. palustris* cells as described (18). Briefly, ∼2 x 10^9^ cells were vacuum-filtered through a nylon membrane (0.45 µm). The membrane was transferred, cell-side down, into a Petri dish containing 2.5 ml of 40:40:20 v/v/v acetonitrile/methanol/water at -20°C and incubated at -20°C for 20 min. The solution was then transferred to a microcentrifuge tube and cells were pelleted at 16,000 x *g* at 4°C for 5 min. Supernatants were transferred to 15 ml-conical tubes and stored at -20°C. Two additional extractions were then applied to the same membrane and combined in the same conical tube before lyophilization and storage at -20°C.

### Liquid chromatography-tandem mass spectrometry (LC-MS/MS)

Compounds were quantified using an Agilent 1290 Infinity II UHPLC coupled to an AB Sciex Qtrap 4000 at the Indiana University (IU) Mass Spectrometry Facility. Analytes were separated on a Waters BEH Amide column (2.1 x 150 mm, 2.5 µm particles) in HILIC mode. Dried samples were reconstituted in 53% mobile phase A plus 47% mobile phase B. Mobile phase A was 95% water, 5% acetonitrile, 20 mM NH_4_OH, and 20 mM ammonium acetate with 5 µM medronic acid. Mobile phase B was 86% acetonitrile, 14% water, 20 mM NH_4_OH and 5 µM medronic acid. The gradient program (flow rate 0.3 ml/min) was: 100% B, 0 to 3 min; ramp 100% to 55% B, 3 and 8 min; hold at 55% B, 8 to 12 min; ramp to 100% B, 12 to 13 min; hold until 36 min. The program was applied for both positive and negative ion modes. QTrap 4000 was operated in multiple reaction monitoring (MRM) mode using Analyst 1.7.1 software. Analyte concentrations were quantified using external calibration curves (Fig S1). Unlabeled standards (purity ≥95%; all from Sigma-Aldrich, except for XMP from Santa Cruz Biotechnology and c-di-GMP from InvivoGen) were diluted in the same solvent as samples for calibration curves.

### Auxotroph bioassays

*R. palustris* cultures (10 ml) were centrifuged in 15-ml conical tubes at 2,415 x *g* for 8 min. Supernatants (3 ml) were injected through a 0.22 μm syringe filter into sterile, sealed, argon (Ar)-filled test tubes. Tubes were then flushed for 0.5 min with Ar and supplemented with 25 mM glucose, 10 mM NH_4_Cl and 1% v/v cation solution. These tubes were then inoculated with washed *E. coli* Δ*purH* or *E. coli* Δ*pyrC* were then inoculated to ∼0.003 OD_660_ and incubated for 16-18 h in the dark.

### Lysate preparation

Cell pellets from 10 ml cultures were resuspended in 0.7 ml of MDC and transferred to 2-ml screw-cap microcentrifuge tubes containing ∼0.25 ml of 0.1 mm Zirconia/Silica beads (BioSpec Products). Cells were lysed in a 4°C room by 8-rounds of bead-beating using a FastPrep^®^-24 homogenizer (MP Biomedical) at max speed for 40 s per round. Lysates were centrifuged at 18,400 x *g* for 20 min at 4°C. Lysate supernatants were then mixed with MDC and prepared as described for bioassays.

### Quantification of live and dead cells by flow cytometry

The Live/dead BacLight Bacterial Viability Kit (Invitrogen) was used according to manufacturer’s instructions with stationary phase *R. palustris* monocultures that were washed and resuspended in 25 mM HEPES (pH 7.5) to a cell density of ∼10^6^ cells/ml. Samples were injected into a NovoCyte flow cytometer (Agilent) with flow rate at 14 μl/min, excited at 488 nm, and emissions detected at 530/30 and 675/30 nm. Populations were analyzed using NovoExpress software (Agilent). Live cell populations were estimated using linear regression of live and dead cell mixtures. Dead cells controls were prepared by incubating with 70% v/v isopropanol for 1 h, then washed and resuspended in HEPES buffer.

### RNA purification

Cultures were grown anaerobically in MDC with 20 mM acetate and 10 mM NH_4_Cl to 0.4-0.8 OD_660_. Cultures were then chilled on ice, transferred to 15-ml conical tubes, and pelleted by centrifugation at 2415 *x g* for 8 min at 4°C. Supernatants were discarded and cell pellets were frozen using dry ice and stored at -80°C. Cells were lysed and RNA extracted using the RNeasy Mini Kit (Qiagen) as per the manufacturer’s instructions, except the lysis step included bead beating as described above. RNA was quantified using a Nanodrop 1000 (Thermo Scientific) at the IU Physical Biochemistry Instrumentation Facility. RNA (20-25 μg) was treated with 4 U Turbo DNase (Ambion) in a 100 μl reaction at 37ºC for 30 min. RNA was then cleaned using the RNeasy MinElute Cleanup Kit (QIAGEN), quantified as before, and adjusted with RNase-free water to 100-200 ng/μl.

### RNA sequencing

RNA (4 μg per sample) was processed by the IU Center for Genomics and Bioinformatics. rRNA was depleted using an Illumina Ribo-Zero Plus rRNA depletion kit. Libraries were prepared using an Illumina TruSeq Stranded mRNA HT kit. Sequencing was performed using an Illumina NextSeq 75-cycle, high-output run. A total of 27-31 million reads were obtained for each sample.

Analysis of differentially expressed genes was performed as described (19, 20) with minor modifications. Briefly, raw reads were preprocessed using Trim Galore v.0.6.6 (https://github.com/FelixKrueger/TrimGalore#readme), a Perl script employing Cutadapt v.1.18 (21), and FastQC v.0.11.5 (22) for trimming of adapter sequences, and removal of low-quality base calls, and quality control, respectively. Processed reads from both CGA0092 and TIE-1 were aligned to the *R. palustris* CGA009 reference genome (NCBI accession#: NC_005296) by HISAT2 v.2.1.0 with options -p, -dta, -no-spliced-alignment, and --rna-strandness RF (23). Samtools v.1.15.1 was used to convert the SAM files output from HISAT2 into BAM format. Aligned reads in BAM format were annotated and the transcript abundance was estimated using StringTie v.1.3.3b (19). Transcript abundance tables were moved into R and DESeq2 was used for differential gene expression analysis (24). Genes unique to either strain could not be considered. Only genes with at least one sample containing ≥ 10 estimated transcript counts were included. Differentially expressed genes with an adjusted *p*-value < 0.05 and a |log2(fold-change)| > 2.0 were considered significant. *Raw reads will be made available at NCBI when the manuscript is accepted*.

### Reverse transcription quantitative real-time PCR (RT-qPCR)

cDNA was prepared from 2 μg RNA using random hexamer primers and SuperScript IV Reverse Transcriptase (RT; Invitrogen) following the manufacturer’s instructions. Standard curves were generated using gDNA. Each gDNA, cDNA, and RT-minus and no template control sample was mixed with 300 nM each forward and reverse primers and 1X SsoAdvanced Universal SYBR Green supermix (Bio-Rad) in a 96-well PCR plate (Eppendorf) for a total volume of 0.1 ml. The thermocycler program was 98°C for 2 min then 40 cycles of 98°C,15 sec; 62°C, 40 sec; 72°C, 30 sec. The reaction was monitored using a Mastercycler ep *realplex* Real-time PCR System (Eppendorf). Data was analyzed by *realplex* software using Noiseband. Primer efficiencies were 94-100%. Specificities were validated by melt curves and by the presence of a single band on a 1% agarose gel.

### Computational modeling

Diffusion of adenine was assessed by modifying a Monod model describing NH_4_^+^ cross-feeding cocultures (7, 25, 26). The model was simplified by omitting H_2_, CO_2_, and ethanol, which do not significantly impact cross-feeding (7, 25, 26). Equations describing N_2_ and NH_4_^+^ were omitted to reflect experimental conditions with saturating NH_4_^+^. The adenine diffusion rate was a product of the *R. palustris* population size, *R. palustris* surface area (27), the adenine permeability coefficient (28), and intracellular adenine concentration. The *E. coli* half-saturation constant for adenine was based on the average Km values for PurP and YicO transporters (29). The model runs in R-studio and is available, along with default parameters, in the supplementary materials.

## Results

### Wild-type *R. palustris* CGA0092 supports *E. coli* purine auxotroph growth

Previously, we found that NH_4_^+^-excreting *R. palustris* Nx supported *E. coli* Δ*purK* purine auxotroph growth in coculture with N_2_ as the sole nitrogen source (8). This result suggested that *R. palustris* Nx externalized purine(s) (Fig 1A). However, Δ*purK* mutants were prone to suppressor mutations, which would complicate our experiments. Therefore, we made an *E. coli* Δ*purH* mutant, which should be less prone to suppression because PurH catalyzes two purine biosynthesis steps (Fig 1B). We verified that *E. coli* Δ*purH* was a purine auxotroph (Fig S2); adenine and adenosine, but not ATP, supported growth. *E. coli* Δ*purH* was also rescued in coculture with *R. palustris* Nx (Fig 1C). We did not observe Δ*purH* suppressors during this study.

In previous cocultures, wild type (WT) *E. coli* was dependent on *R. palustris* Nx for NH_4_^+^ (7) (Fig 1A, top). Adding NH_4_Cl decoupled the growth rates of the partners and led to coculture growth trends resembling an *E. coli* monoculture (7) (Fig 1C). We wondered if cocultures with *E. coli* Δ*purH* would respond similarly to NH_4_Cl or if the growth rate would be limited by purine availability (Fig 1A, bottom). Indeed, growth trends for NH_4_Cl-supplied *R. palustris* Nx + *E. coli* Δ*purH* cocultures more closely resembled NH_4_^+^-cross-feeding cocultures than NH_4_Cl-supplied cocultures with WT *E. coli* (Fig 1C).

Purine externalization seemed costly, so we questioned whether it was due to engineered mutations. We thus attempted to coculture *E. coli* Δ*purH* with WT *R. palustris* CGA0092. This strain also supported *E. coli* Δ*purH* in coculture (Fig 1D). Thus, purine externalization is not a result of any engineered mutations.

### Purine externalization is not dependent on *E. coli* or growth conditions

In some cross-feeding systems, the recipient can influence metabolite externalization by a producer (30-32). To test whether *E. coli* induces *R. palustris* purine externalization, we inoculated *E. coli* Δ*purH* into media supplemented with *R. palustris* monoculture supernatant. *E. coli* Δ*purH* grew proportionately to the amount of supernatant supplied (Fig S3) and to the *R. palustris* population size that generated the supernatant, regardless of the *R. palustris* growth phase (Fig 1E). Thus, *R. palustris* purine externalization occurs during exponential growth and is not induced by *E. coli*.

Thus far, purine externalization was only observed under photoheterotrophic conditions, where energy is derived from light and carbon from organic compounds. However, *R. palustris* can grow in diverse conditions that we hypothesized could affect purine externalization. We thus examined *E. coli* Δ*purH* growth with CGA0092 supernatants from photoautotrophic conditions, and from chemotrophic conditions requiring aerobic or anaerobic respiration. *E. coli* Δ*purH* growth, a proxy for purine externalization, was roughly correlated with the amount of *R. palustris* growth rather than growth condition (Fig 1F) or growth rate (Fig S4).

### Purine cross-feeding supports coexistence

Cocultures are most useful as experimental systems if they support stable coexistence. To assess if purine cross-feeding supports coexistence, we performed an invasion-from-rare assay (17). We used a biofilm-deficient *R. palustris* strain (CGA4004) to improve population measurement accuracy by CFUs (CGA4004 gives similar coculture growth trends as CGA0092, Fig 1D). *E. coli* Δ*purH* exhibited negative frequency-dependent selection (Fig 2A). The x-intercept suggests an *E. coli* Δ*purH* equilibrium frequency of ∼ 2.3 % in coculture. Indeed, in cocultures where we tracked populations over time by CFUs, the *E. coli* Δ*purH* frequency moved from 68% to 3-5% (Fig 2B). This *E. coli* frequency is similar to those observed when *E. coli* and *R. palustris* growth rates are coupled by NH_4_^+^ cross-feeding (7, 25, 26, 30, 33).

**Fig 2.**
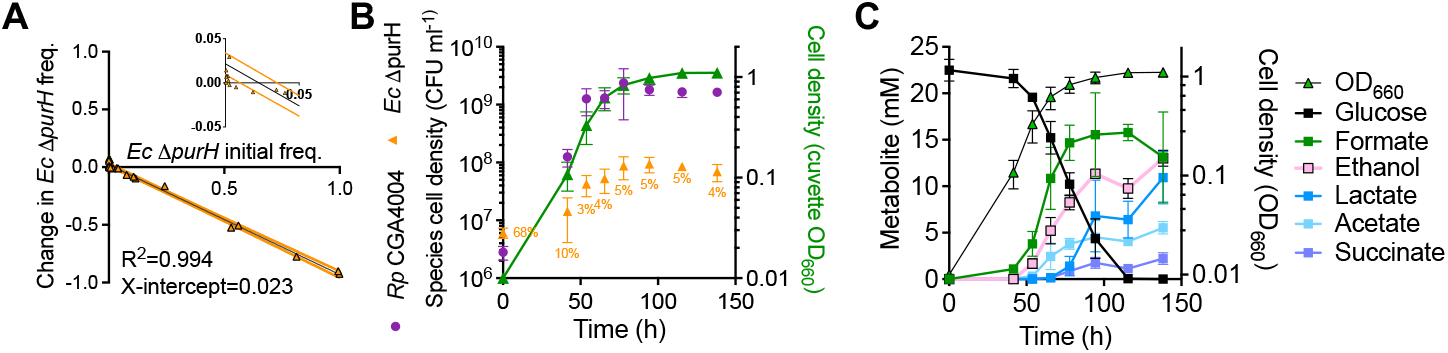
Purine cross-feeding supports coexistence. A. Invasion-from-rare assay pairing *E. coli* Δ*purH* with CGA4004 in cocultures. Linear regression (black line) +/- 95% confidence intervals (orange lines) was applied. The inset graph is an enlarged portion of the same data to help visualize the x-intercept. **B, C**. Population trends (**B**) and metabolic trends (**C**) in batch *R. palustris* CGA4004 + *E. coli* Δ*purH* cocultures. Percentages refer to the *E. coli* frequency. Error bars = SD; n=3. Some error bars are smaller than the symbols.

We also questioned how purine cross-feeding affected organic acid cross-feeding. When glucose and fermentation products were tracked by HPLC we saw an accumulation of organic acids that *R. palustris* can consume (consumable organic acids) accumulated (Fig 2C). This accumulation suggests that CGA4004 + *E. coli* Δ*purH* cocultures are metabolically similar to NH_4_^+^-cross-feeding cocultures that used an *R. palustris* with a 3-fold high NH_4_^+^ excretion rate than the NifA* strain, causing *E. coli* to excrete organic acids faster than *R. palustris* could consume them (7).

### *R. palustris* TIE-1 does not support *E. coli* Δ*purH* growth

*R. palustris* TIE-1 (34) is closely related to CGA0092; the two strains have identical 16S rRNA sequences and share 5.28 Mb of DNA with 97.9% identity (35). We tested whether TIE-1 also externalizes purines by attemtping to coculture it with *E. coli* Δ*purH*. These cocultures exhibited linear growth (Fig 3A), suggesting that TIE-1 grew on organic acids released by non-growing *E. coli*, a phenomenon we characterized previously in nitrogen-starved cocultures (26). Indeed, *E. coli* Δ*purH* populations declined in coculture with TIE-1 but increased in coculture with CGA0092 (Fig 3B). We verified that the results were not influenced by different growth traits; monoculture growth curves for the two strains were similar (Fig S4). We also confirmed that the decline in *E. coli* Δ*purH* populations was not due to inhibitory factors produced by TIE-1; *E. coli* Δ*purH* grew in TIE-1 monoculture supernatants when supplemented with adenine (Fig 3C). Thus, TIE-1 does not externalize enough purine(s) to support *E. coli* Δ*purH* growth.

**Fig 3.**
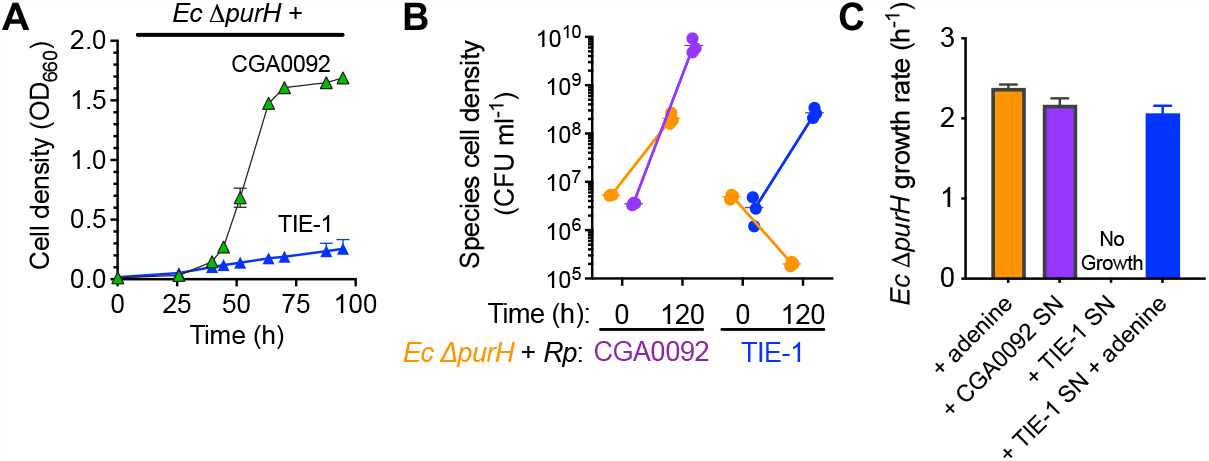
*R. palustris (Rp)* TIE-1 does not support *E. coli (Ec)* Δ*purH* auxotroph growth. **A.** Coculture growth curves of *E. coli* Δ*purH* with *R. palustris* CGA0092 or TIE-1. **B**. Comparison of initial and final population sizes by colony forming units (CFUs) in cocultures pairing *E. coli* Δ*purH* with CGA0092 or TIE-1. **C**. Growth of *E. coli* Δ*purH* +/-CGA0092 or TIE-1 supernatants +/- 50 μM adenine. **A-C**. Error bars = SD; n=3. SN, supernatant.

### *R. palustris* CGA0092 externalizes adenine

We sought to identify the purine(s) externalized by CGA0092. Taking advantage of the lack of purine externalization by TIE-1, we used LC-MS/MS to compare nucleobase-containing compounds in between the two strains in monocultures. For each strain, similar concentrations were observed between exponential phase and stationary phase for both intracellular and extracellular samples, with the exception of adenine (Fig 4A, B; Table S4, S5). Adenine was measured at 17 ± 2 μM/OD_660_ in CGA0092 exponential phase supernatants, which was 57-fold higher than in TIE-1 supernatants (0.3 ± 0.1 μM) (Fig 4A). Intracellular adenine concentrations were 89-fold higher in CGA0092 (1.52 ± 0.64 mM) compared to TIE-1 (0.02 ± 0.00 mM) (Fig 4B).

**Fig 4.**
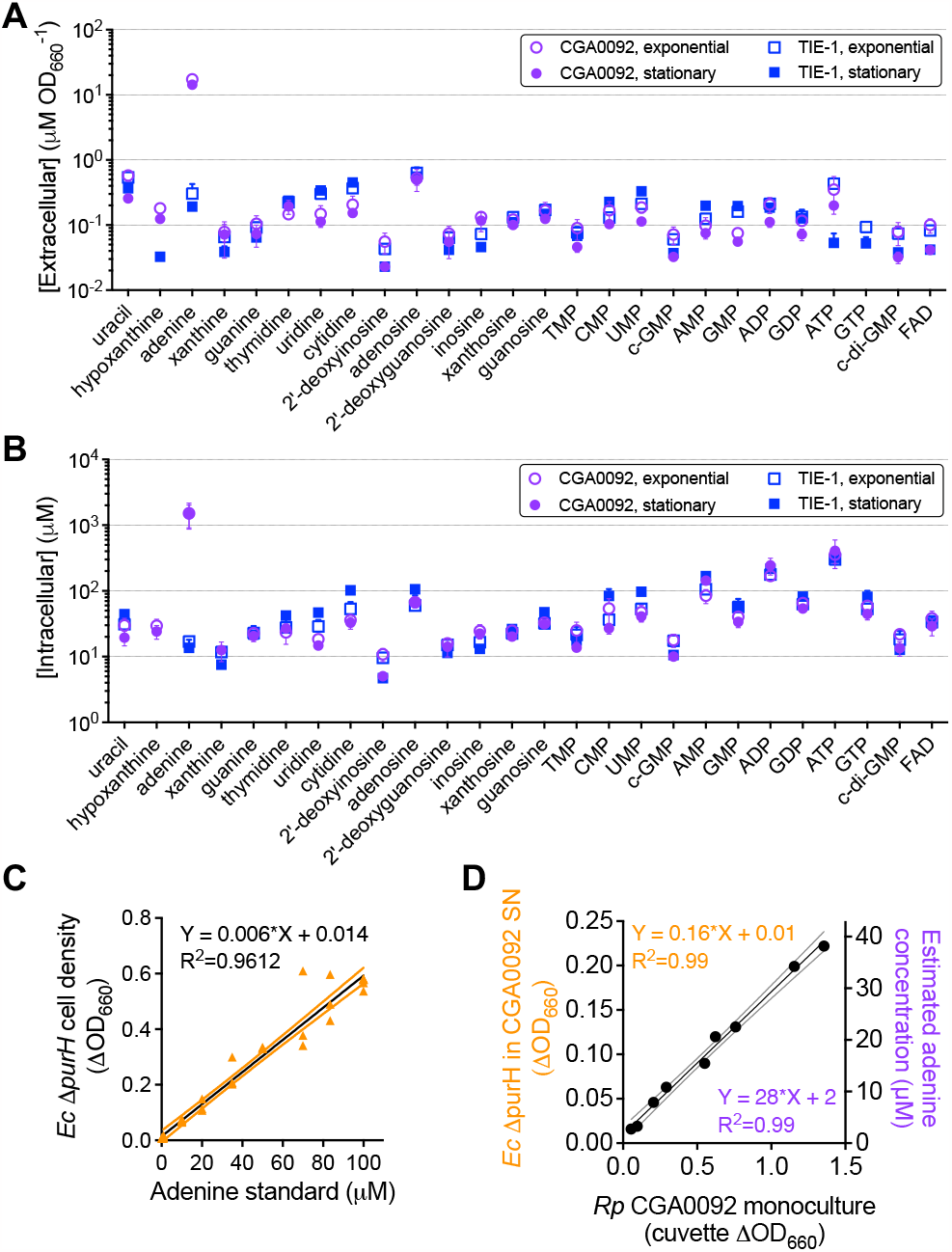
*R. palustris* CGA0092 externalizes adenine. A, B. Extracellular (**A**) and intracellular (**B**) concentrations of nucleobase-containing compounds from monocultures of *R. palustris* CGA0092 and TIE-1. Molecules are arranged by increasing molecular weight. **C**. Standard curve for quantifying adenine in CGA0092 supernatants (or bioavailable purines in general) using a *E. coli* Δ*purH* bioassay. **D**. Estimated extracellular adenine in CGA0092 monoculture supernatants using the *E. coli* Δ*purH* bioassay. **C, D**. Outer lines for each linear regression analysis represent the 95% confidence interval. SN, supernatant.

We then developed a bioassay to facilitate quantification of external adenine. Using a standard curve of *E. coli* Δ*purH* cell density versus adenine concentration (Fig 4C), we estimated adenine levels of 28 +/- 2 μM /CGA0092 OD_660_ (Fig 4D),which was 1.6-fold higher than that determined by LC-MS/MS. Although the bioassay is responsive to other purines (Fig S2), the discrepancy likely stems from differences in methodology. Aside from adenine, LC-MS/MS indicated that other purine levels were similar between CGA0092 and TIE-1 (Fig 4A). Thus, if the other purines accounted for the discrepancy, we would expect *E. coli* Δ*purH* to grow 1.6-fold higher in CGA0092 supernatants than in TIE-1 supernatants, but *E. coli* Δ*purH* does not grow in TIE-1 supernatants (Fig 3C). Although it is possible that LC-MS/MS overlooked some purines, we can conclude that *E. coli* Δ*purH* growth in CGA0092 supernatants is primarily due to adenine.

### Adenine externalization can be explained by diffusion across the membrane

Having identified adenine accumulation in CGA0092, we next pursued how adenine is externalized. We first addressed lysis by comparing live and dead cells frequencies in CGA0092 versus TIE-1 monocultures. Both strains shared a similarly low frequency of dead cells, suggesting that lysis is not a major contributor to CGA0092 adenine externalization (Fig 5A). A lack of adenine externalization due to lysis was also supported by experiments in which we grew *E. coli* Δ*purH* and a pyrimidine auxotroph control (Δ*pyrC*) with CGA0092 cell lysates. CGA0092 cell lysate only supported ∼25% of the *E. coli* Δ*purH* growth observed in supernatants from an equivalent number of cells (Fig 5B). Lysate also supported *E. coli* Δ*pyrC* growth, indicating that supernatants are specifically enriched in purines (i.e., adenine) whereas lysate is not (Fig 5B).

**Fig 5.**
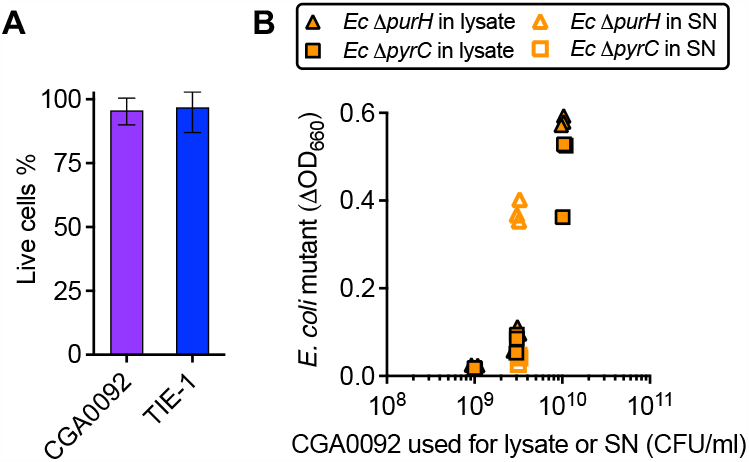
Cell lysis cannot explain *E. coli* Δ*purH* growth in CGA0092 supernatants. A. Live-dead stains of stationary phase CGA0092 and TIE-1 monocultures. Approximately 3 x 10^7^ and 5 x 10^7^ cells were counted for CGA0092 and TIE-1, respectively. Error bars, SD. **B**. Comparison of *E. coli* Δ*purH* (purine auxotroph) and Δ*pyrC* (pyrimidine auxotroph) growth in CGA0092 supernatants (open symbols) and lysates (closed symbols). Each symbol represents a biological replicate (n=3). SN, supernatant.

With lysis ruled out, we addressed diffusion across the cytoplasmic membrane. We modified a Monod model that simulated coculture trends based on user-specified NH_4_^+^ excretion levels to instead simulate adenine externalization as a function of an adenine permeability coefficient (28), *R. palustris* surface area (27), and intracellular adenine concentration (Fig 4B). The model accurately predicted extracellular adenine levels for both CGA0092 and TIE-1 monocultures using both default parameters and for a range of realistic cell sizes (27), a two-fold difference in intracellular adenine, and a > 2-fold change in permeability (Fig 6A, B).

**Fig 6.**
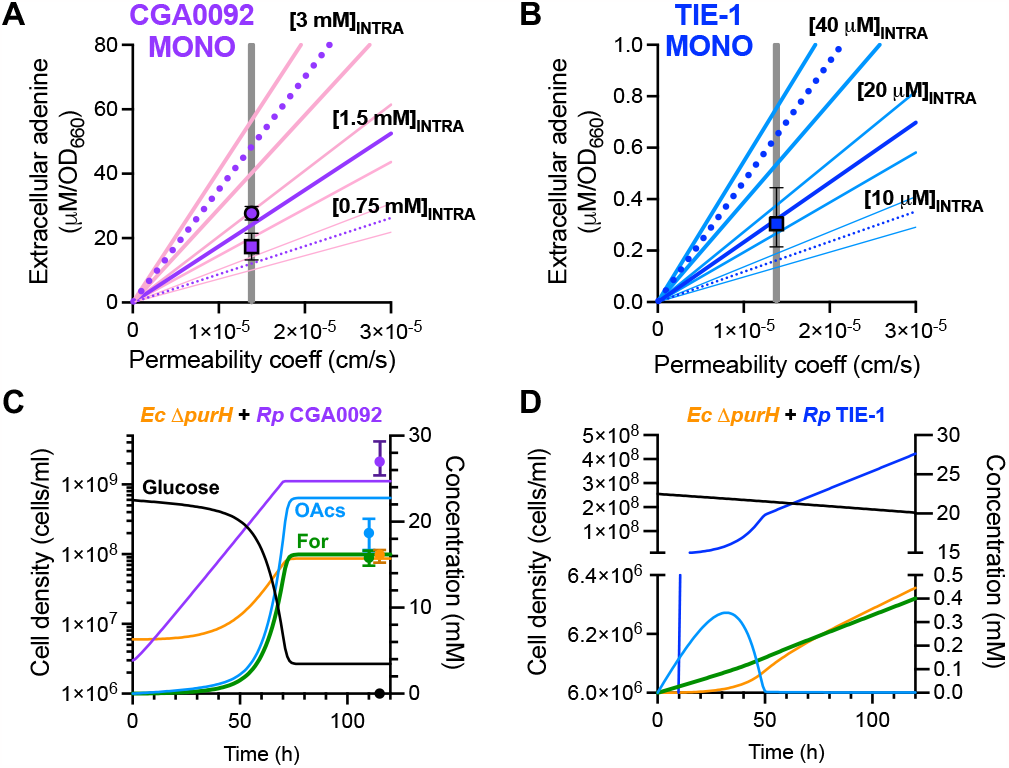
Simulations using adenine diffusion across a membrane accurately predict extracellular adenine levels. Adenine externalization was simulated as a function of the permeability coefficient, cell surface area, and intracellular concentration using a Monod model (supplementary information). **A, B**. Monoculture simulations were run for three different intracellular adenine concentrations (purple (A) or dark blue (B) lines) and three different cell sizes (pink (A) or light blue (B) lines represent the upper and lower bounds on cell size (27); supplementary information) across a range of adenine permeability coefficients (coeff). Gray line, published adenine permeability coefficient (28); square symbol, extracellular adenine measured by LC-MS/MS; round symbol, extracellular adenine measured by a bioassay. Symbols were arbitrarily placed at the published adenine permeability coefficient. Error bars = SD; n=3. **C, D**. Coculture simulations using the published adenine permeability coefficient, average cell size, intracellular adenine concentrations measured by LC-MS/MS (supplementary information. **C**. Symbols represent final empirical values, arbitrarily placed to avoid overlap with the y-axis. Error bars = range; n=3.

The model also accurately predicted the final *E. coli* population in coculture with CGA0092 (Fig 6C). The model fell short of predicting observed *R. palustris* populations because it was over-sensitive to the inhibitory effect of pH, limiting the *R. palustris* population size, glucose consumption, and organic acid consumption. However, when we took acid inhibition out of the model, the predicted *E. coli* population was still within the observed range (not shown). We did not attempt to make quantitative predictions with TIE-1 cocultures because our model does not describe *E. coli* death, which occurs (Fig 3B). However, the model still accurately predicted that the level of adenine externalization by TIE-1 cannot support substantial *E. coli* growth, resulting in linear TIE-1 population growth (Fig. 6D). Thus, adenine excretion can likely be explained by diffusion across a membrane, a literal definition of leakage (2). For this reason, we did not pursue possible efflux proteins, though we cannot rule out their involvement.

### CGA0092 adenine externalization is likely influenced by low *apt* expression

To address why CGA0092 accumulates adenine, we again exploited a CGA0092 versus TIE-1 comparison. Given their high genetic relatedness, we reasoned that RNAseq could reveal insightful gene expression differences. Indeed, of the 4,757 genes compared, only 581 showed significantly different transcript levels; 64 genes had higher transcript levels in TIE-1 and 517 were lower (Supplementary materials). Of primary interest was the relatively low CGA0092 transcript levels for *apt* (TX73_RS22960, RPA4492), encoding the purine salvage enzyme adenine phosphoribosyltransferase (Apt). RT-qPCR analysis verified the difference in *apt* gene expression (Fig 7A).

**Fig. 7.**
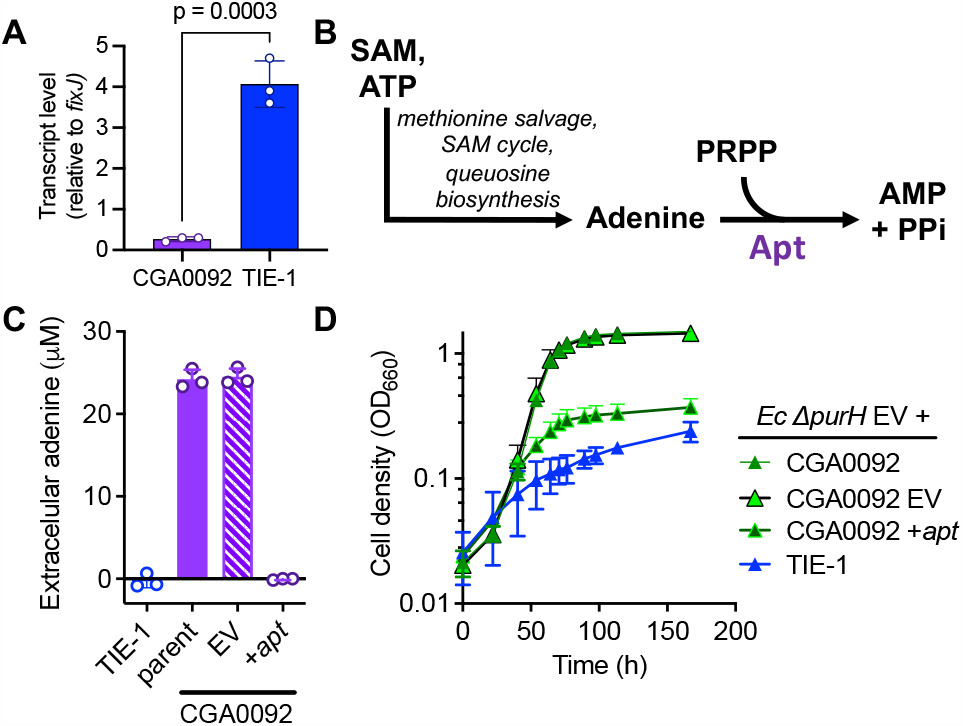
Ectopic expression of Apt decreases CGA0092 adenine externalization. **A.** Transcript levels of *apt* in CGA0092 and TIE-1 relative to the house-keeping gene *fixJ* determined by RT-qPCR. **B**. Adenine salvage pathway, showing the location of the Apt enzyme. **C**. Extracellular purines (assumed to be adenine) in *R. palustris* supernatants with and without ectopic *apt* expression. Adenine was measured using the *E. coli* Δ*purH* bioassay. **D**. *R. palustris* + *E. coli* Δ*purH* coculture growth curves with and without ectopic *apt* expression. **C, D**. Error bars = SD; n=3. EV, empty vector pBBPgdh; +*apt*, constitutive expression vector pBBPgdh-*apt*.

We hypothesized that low CGA0092 *apt* expression creates a bottleneck, leading to adenine accumulation and leakage (Fig 7B). If so, higher expression should alleviate the bottleneck. Thus, we expressed CGA0092 *apt* under a constitutive promoter from a plasmid and measured extracellular adenine using the bioassay. Supernatants from CGA0092 with and without an empty vector had similar adenine levels, whereas adenine was undetectable in supernatants from CGA0092 expressing *apt* from a plasmid (Fig 7C). Similar trends were seen in cocultures with *E. coli* Δ*purH*; the CGA0092 empty vector control gave coculture growth curves similar to those with CGA0092 whereas CGA0092 expressing *apt* from a plasmid resulted in poor coculture growth (Fig. 7D).

### Orientation of *apt* does not always indicate purine externalization

We questioned why *apt* expression is low in CGA0092 compared to TIE-1. The *apt* genes, plus 73 nucleotides upstream (the entire intergenic region for TIE-1) are identical in the two strains. However, *apt* is part of a gene cluster that has an opposite orientation in CGA0092 versus TIE-1 (Fig 8 inset). We hypothesized that gene orientation affects *apt* expression and adenine externalization. This hypothesis is supported by RNAseq data showing that *lemA*, on the other side of the cluster, had 2.2-fold higher transcript levels in CGA0092 (Fig S6 and Supplementary materials), suggesting possible read-through from *trpE*.

**Fig 8.**
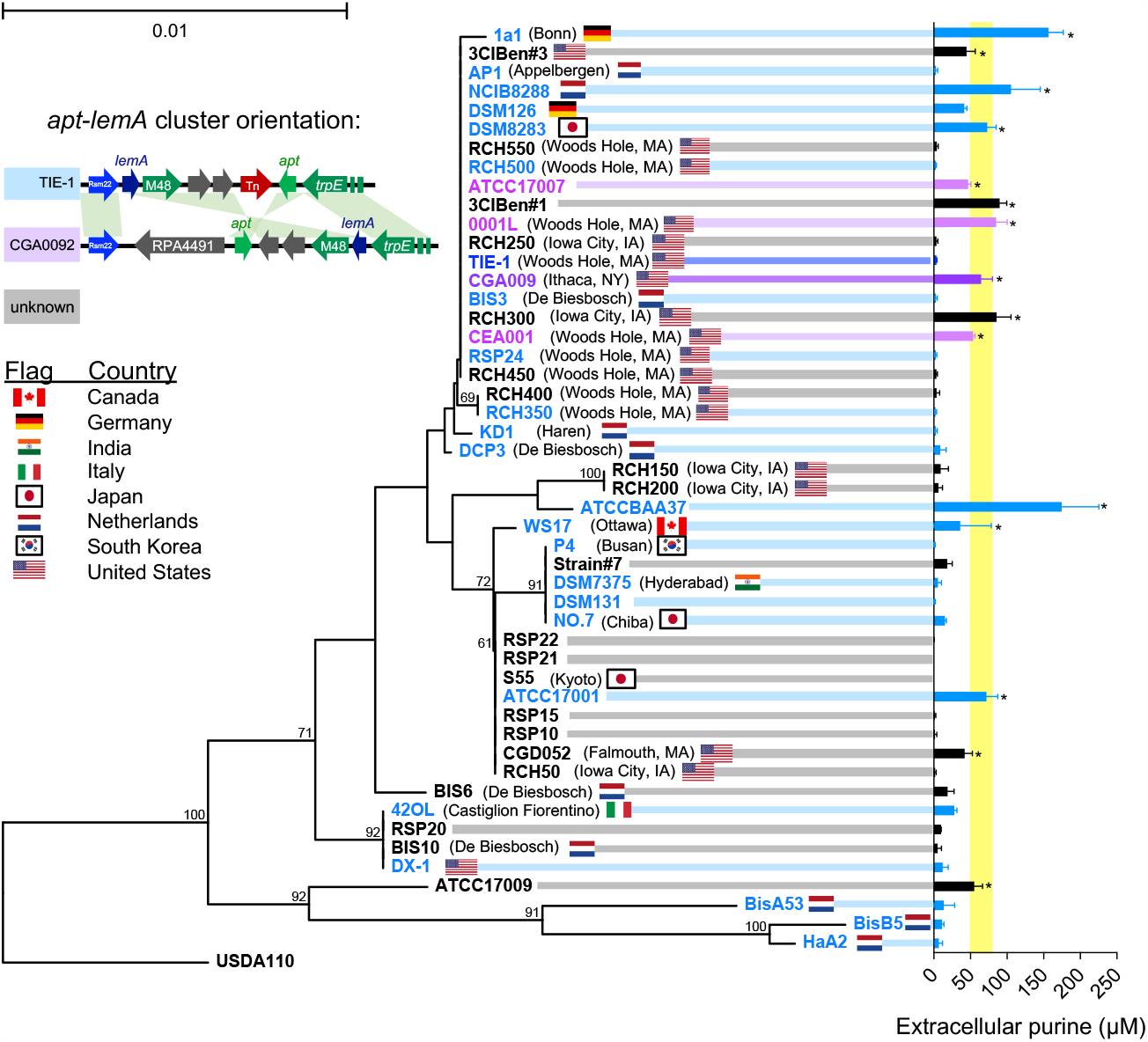
Diverse R. palustris strains excrete purine, regardless of apt orientation. Phylogenetic relationships of 49 R. *palustris* strains tested for adenine excretion based on 16S rRNA sequences. Bracketed text and flags indicate isolation location where known. Bootstrap values (100 replicates) are at branch points (only showing values > 50). The bar represents substitutions per site. *Bradyrhizobium diazoefficiens* USDA110 was used to root the tree. Purple or blue tree shading indicate CGA009 or TIE-1 *apt* gene orientation, respectively. Gray shading indicates unknown *apt* gene orientation. **Right**. Extracellular purine measured using the *E. coli* Δ*purH* bioassay. Yellow shading indicates the CGA0092 standard deviation. ^*^, significantly more purine than TIE-1 from One-way ANOVA with a Dunnett correction for multiple comparisons; p < 0.1. Error bars = SD, n=3. **Inset**. CGA0092 and TIE-1 *apt-lemA* clusters shown to scale. Other bacteria with the TIE-1 orientation do not have the transposon (Tn; Fig S7).

To explore whether *apt* orientation can explain adenine externalization, we used the bioassay to measure extracellular adenine (and possibly other purines) from 49 *R. palustris* strains (Fig 8). Fifteen strains externalized significantly more purine than TIE-1 (Fig 8). The CGA0092-like *apt* orientation was always associated with purine externalization. Overall, the Spearman correlation coefficient for purine externalization and *apt* orientation was significant (p = 0.026). Even so, some strains with TIE-1-like orientation also externalized purine. Thus, other factors in additional to *apt* orientation can contribute to purine externalization. This experiment also revealed that purine externalization is relatively common among *R. palustris* isolates.

## Discussion

We determined that WT *R. palustris* CGA0092 externalizes adenine in quantities that can sustain an *E. coli* purine auxotroph. Based on computational modeling, RNAseq, and mutational approaches, we propose that adenine accumulates intracellularly due to a bottleneck in adenine salvaging, and then leaves the cell by diffusion across the cytoplasmic membrane. Purine externalization was also observed in several other *R. palustris* isolates, suggesting that this phenomenon could occur in nature (Fig 8). Purine excretion by *R. palustris* is yet another example of cross-feeding of communally valuable metabolites, joining other examples that include amino acids (36-38) and vitamins (31, 38-40).

### What is the role of *R. palustris* adenine externalization?

*R. palustris* strains from diverse regions exhibited purine externalization, suggesting that it is not an artifact of domestication and that it could play a physiological, signaling, or ecological role. Adenine externalization could be analogous to uracil externalization by *E. coli*, which helps maintain high growth rates in response to perturbations to intracellular pyrimidine pools (41). For CGA0092, adenine externalization is a constant feature of exponential growth across diverse conditions (Fig 1E, F). Thus, if adenine externalization maintains homeostatic metabolite levels, it is both constitutive and effective; aside from adenine, intracellular nucleobase-containing metabolite levels were similar to those in TIE-1 (Fig 4B).

Adenine externalization could also be due to overflow metabolism caused by a level of carbon influx not normally experienced in nature (42, 43). If true, one might expect a correlation between growth rate and adenine externalization (44). However, we did not observe a strong correlation with growth rate, at least when growth rate was determined by the growth condition (Fig S4). The observation that some closely related strains do not externalize adenine (Fig 8), despite having similar growth trends (Fig S5), also argues against overflow metabolism due to carbon availability.

Extracellular adenine could also participate in signaling and/or cross-feeding. Some pathogens externalize ATP as a signal molecule for its immunosuppressing effects (45). *R. palustris* has features that suggest interbacterial and interkingdom relationships. Similar to its reliance on other organisms to convert sugars into organic acids, *R. palustris* relies on lignolytic organisms to release consumable lignin monomers (46). Lignin also plays into *R. palustris* quorum sensing, a cell-density dependent intercellular signaling mechanism. CGA009 can only make the p-coumaroyl homoserine lactone signal when supplied with lignin monomers, hinting at a broader relationship with plants or lignolytic fungi (47). Perhaps purine externalization by diverse *R. palustris* isolates also hints at an inter-organismal relationship (Fig 8).

### Mechanisms of metabolite externalization

Our study is one of the few to address how cross-fed metabolites are externalized. While it is often assumed that metabolites passively leak from bacteria, transporters are likely involved for most charged and polar molecules (2). Some purine efflux proteins are also known (48, 49). However, our model suggests that adenine, an uncharged molecule and the most hydrophobic of all the molecules that we measured by LC-MS/MS, can escape via diffusion across the cytoplasmic membrane. Our results show that if the pool of intracellular adenine is large enough, adenine leakage can lead to cross-feeding. If a nearby microbe can reciprocate, and close proximity can be maintained, then cross-feeding could become subject to selection (3, 4, 6).

Although diffusion can explain adenine externalization, we cannot rule out involvement of efflux proteins. The involvement of efflux proteins might be suggested by the similar adenine concentrations observed between exponential and stationary phase; adenine externalization halted with growth (Fig. 4). However, similar levels could also be explained if leakage and uptake reached a steady state in stationary phase, or if changes in membrane composition during stationary phase, like the generation of cyclopropane fatty acids (50), limited adenine leakage.

### Elusive signatures for metabolite externalization

Predictions of cross-feeding interactions from genomic data are often based on the absence of biosynthetic genes (5, 51). Such missing genes are suggestive, but not concrete, indicators that an organism competitively or cooperatively acquires essential metabolites from a neighbor. Moreover, when considering externalization of communally valuable metabolites, we lack even suggestive genetic signatures.

We hypothesized that *apt* orientation was a signature for adenine externalization. Whereas the orientation observed in CGA0092 was always correlated with purine externalization, all strains with this orientation were also closely related (Fig 8). Some strains that had the opposite *apt* orientation also externalized purine(s), though we do not currently know if the purine was adenine in these cases. Thus, *apt* orientation might be one driver of adenine externalization but other unknown factors can also be sufficient. Even if *apt* orientation is a signature for adenine externalization, the synteny of the reversible gene cluster is not conserved outside of *R. palustris*.

Our study thus highlights a challenge in predicting metabolite-externalizing subpopulations. Genomic signatures could be more diverse than the repertoire of cross-fed metabolites. Metabolite externalization can also be conditional (2, 31, 32, 41) and need not even require a genetic signature in the producer. Previously we found that enhanced metabolite acquisition by a recipient was sufficient to stimulate more NH_4_^+^ release by a producer and establish cross-feeding; the genetic signature was in the recipient (30). With such confounding elements, mechanistic studies into microbial interactions will remain essential to obtain a breadth of knowledge on the factors governing metabolite externalization before we can make accurate predictions about cross-feeding from the available wealth of genomic information.

## Acknowledgements

This work was supported in part by the US Army Research Office grants W911NF-14-1-0411 (JBM), W911NF-17-1-0159 (JBM), and W911NF-21-1-0015 (CSH), the National Science Foundation CAREER award MCB-1749489 (JBM), and the NIH grant R35GM150625 (MB). Supercomputing resources were supported in part by Lilly Endowment, Inc., through its support for the IU Pervasive Technology Institute.

We are grateful to A. Dalia, J. Drummond, J.P. Gerdt, D. Rusch, and the McKinlay lab for helpful advice. We also thank C. Dann and C. Bauer for c-di-GMP and c-GMP standards, respectively.

## References

1. Fritts RK, McCully AL, McKinlay JB. 2021. Extracellular metabolism sets the table for microbial cross-feeding. Microbiol Mol Biol Rev 85:e00135–20.

2. McKinlay JB. 2023. Are bacteria leaky? Mechanisms of metabolite externalization in bacterial cross-feeding. Annu Rev Microbiol doi:10.1146/annurev-micro-032521-023815.

3. Pande S, Kost C. 2017. Bacterial unculturability and the formation of intercellular metabolic networks. Trends Microbiol 25:349–361.

4. D’Souza G, Shitut S, Preussger D, Yousif G, Waschina S, Kost C. 2018. Ecology and evolution of metabolic cross-feeding interactions in bacteria. Nat Prod Rep 35:455–488.

5. Zengler K, Zaramela LS. 2018. The social network of microorganisms — how auxotrophies shape complex communities. Nat Rev Microbiol 16:383–390.

6. Momeni B, Chen C-C, Hillesland KL, Waite A, Shou W. 2011. Using artificial systems to explore the ecology and evolution of symbioses. Cell Mol Life Sci 68:1353–1368.

7. LaSarre B, McCully AL, Lennon JT, McKinlay JB. 2017. Microbial mutualism dynamics governed by dose-dependent toxicity of cross-fed nutrients. ISME J 11:337–348.

8. LaSarre B, Deutschbauer AM, Love CE, McKinlay JB. 2020. Covert cross-feeding revealed by genome-wide analysis of fitness determinants in a synthetic bacterial mutualism. Appl Environ Microbiol 86:e00543–20.

9. Fritts RK, LaSarre B, Stoner AM, Posto AL, McKinlay JB. 2017. A Rhizobiales-specific unipolar polysaccharide adhesin contributes to Rhodopseudomonas palustris biofilm formation across diverse photoheterotrophic conditions. Appl Environ Microbiol 83:e03035–16.

10. Blattner FR, Plunkett G, 3rd, Bloch CA, Perna NT, Burland V, Riley M, Collado-Vides J, Glasner JD, Rode CK, Mayhew GF, Gregor J, Davis NW, Kirkpatrick HA, Goeden MA, Rose DJ, Mau B, Shao Y. 1997. The complete genome sequence of Escherichia coli K-12. Science 277:1453–62.

11. Datsenko KA, Wanner BL. 2000. One-step inactivation of chromosomal genes in Escherichia coli K-12 using PCR products. Proc Natl Acad Sci USA 97:6640–6645.

12. Baba T, Ara T, Hasegawa M, Takai Y, Okumura Y, Baba M, Datsenko KA, Tomita M, Wanner BL, Mori H. 2006. Construction of Escherichia coli K-12 in-frame, single-gene knockout mutants: the Keio collection. Mol Syst Biol 2:2006.0008.

13. Kostylev M, Otwell AE, Richardson RE, Suzuki Y. 2015. Cloning should be simple: Escherichia coli DH5α-mediated assembly of multiple DNA fragments with short end homologies. PLoS One 10:e0137466.

14. Pelletier DA, Hurst GB, Foote LJ, Lankford PK, McKeown CK, Lu TY, Schmoyer DD, Shah MB, Hervey WJt, McDonald WH, Hooker BS, Cannon WR, Daly DS, Gilmore JM, Wiley HS, Auberry DL, Wang Y, Larimer FW, Kennel SJ, Doktycz MJ, Morrell-Falvey JL, Owens ET, Buchanan MV. 2008. A general system for studying protein-protein interactions in Gram-negative bacteria. J Proteome Res 7:3319–28.

15. LaSarre B, Morlen R, Neumann GC, Harwood CS, McKinlay JB. 2023. Nitrous oxide reduction by two partial denitrifying bacteria requires denitrification intermediates that cannot be respired. bioRxiv:10.1101/2022.06.21.497020.

16. McKinlay JB, Zeikus JG, Vieille C. 2005. Insights into Actinobacillus succinogenes fermentative metabolism in a chemically defined growth medium. Appl and Environ Microbiol 71:6651–6656.

17. Hammarlund SP, Chacón JM, Harcombe WR. 2019. A shared limiting resource leads to competitive exclusion in a cross-feeding system. Environ Microbiol 21:759–771.

18. Bennett BD, Yuan J, Kimball EH, Rabinowitz JD. 2008. Absolute quantitation of intracellular metabolite concentrations by an isotope ratio-based approach. Nat Protoc 3:1299–311.

19. Pertea M, Kim D, Pertea GM, Leek JT, Salzberg SL. 2016. Transcript-level expression analysis of RNA-seq experiments with HISAT, StringTie and Ballgown. Nat Protoc 11:1650–67.

20. Behringer MG, Ho WC, Meraz JC, Miller SF, Boyer GF, Stone CJ, Andersen M, Lynch M. 2022. Complex ecotype dynamics evolve in response to fluctuating resources. mBio 13:e0346721.

21. Martin M. 2011. Cutadapt removes adapter sequences from high-throughput sequencing reads. EMBnet j 17.

22. Andrews S. 2010. FastQC: A quality control tool for high throughput sequence data. Babraham Bioinformatics https://www.bioinformatics.babraham.ac.uk/projects/fastqc/.

23. Kim D, Paggi JM, Park C, Bennett C, Salzberg SL. 2019. Graph-based genome alignment and genotyping with HISAT2 and HISAT-genotype. Nat Biotechnol 37:907–915.

24. Love MI, Huber W, Anders S. 2014. Moderated estimation of fold change and dispersion for RNA-seq data with DESeq2. Genome Biol 15:550.

25. McCully AL, LaSarre B, McKinlay JB. 2017. Recipient-biased competition for an intracellularly generated cross-fed nutrient is required for coexistence of microbial mutualists. mBio 8:e01620–17.

26. McCully AL, LaSarre B, McKinlay JB. 2017. Growth-independent cross-feeding modifies boundaries for coexistence in a bacterial mutualism. Environ Microbiol 19:3538–3550.

27. LaSarre B, Kysela DT, Stein BD, Ducret A, Brun YV, McKinlay JB. 2018. Restricted localization of photosynthetic intracytoplasmic membranes (ICMs) in multiple genera of purple nonsulfur bacteria. mBio 9:e00780–18.

28. Xiang TX, Anderson BD. 1994. The relationship between permeant size and permeability in lipid bilayer membranes. J Membr Biol 140:111–22.

29. Papakostas K, Botou M, Frillingos S. 2013. Functional identification of the hypoxanthine/guanine transporters YjcD and YgfQ and the adenine transporters PurP and YicO of Escherichia coli K-12. J Biol Chem 288:36827–40.

30. Fritts RK, Bird JT, Behringer MG, Lipzen A, Martin J, Lynch M, McKinlay JB. 2020. Enhanced nutrient uptake is sufficient to drive emergent cross-feeding between bacteria in a synthetic community. ISME J 14:2816–2828.

31. Grant MAA, Kazamia E, Cicuta P, Smith AG. 2014. Direct exchange of vitamin B_12_ is demonstrated by modelling the growth dynamics of algal–bacterial cocultures. ISME J 8:1418–1427.

32. Bunbury F, Deery E, Sayer AP, Bhardwaj V, Harrison EL, Warren MJ, Smith AG. 2022. Exploring the onset of B_12_-based mutualisms using a recently evolved Chlamydomonas auxotroph and B_12_-producing bacteria. Environ Microbiol 24:3134–3147.

33. McCully AL, Behringer MG, Gliessman JR, Pilipenko EV, Mazny JL, Lynch M, Drummond DA, McKinlay JB. 2018. An Escherichia coli nitrogen starvation response is important for mutualistic coexistence with Rhodopseudomonas palustris. Appl Environ Microbiol 84:e00404–18.

34. Jiao Y, Kappler A, Croal LR, Newman DK. 2005. Isolation and characterization of a genetically tractable photoautotrophic Fe(II)-oxidizing bacterium, Rhodopseudomonas palustris strain TIE-1. Appl Environ Microbiol 71:4487–96.

35. Oda Y, Larimer FW, Chain PSG, Malfatti S, Shin MV, Vergez LM, Hauser L, Land ML, Braatsch S, Beatty JT, Pelletier DA, Schaefer AL, Harwood CS. 2008. Multiple genome sequences reveal adaptations of a phototrophic bacterium to sediment microenvironments. Proc Natl Acad Sci USA 105:18543–18548.

36. Mee MT, Collins JJ, Church GM, Wang HH. 2014. Syntrophic exchange in synthetic microbial communities. Proc Natl Acad Sci U S A 111:E2149–56.

37. Wintermute EH, Silver PA. 2010. Emergent cooperation in microbial metabolism. Mol Syst Biol 6:407.

38. Martinson JNV, Chacón JM, Smith BA, Villarreal AR, Hunter RC, Harcombe WR. 2023. Mutualism reduces the severity of gene disruptions in predictable ways across microbial communities. bioRxiv:10.1101/2023.05.08.539835.

39. Croft MT, Lawrence AD, Raux-Deery E, Warren MJ, Smith AG. 2005. Algae acquire vitamin B_12_through a symbiotic relationship with bacteria. Nature 438:90–93.

40. Kazamia E, Czesnick H, Nguyen TT, Croft MT, Sherwood E, Sasso S, Hodson SJ, Warren MJ, Smith AG. 2012. Mutualistic interactions between vitamin B_12_-dependent algae and heterotrophic bacteria exhibit regulation. Environ Microbiol 14:1466–76.

41. Reaves ML, Young BD, Hosios AM, Xu YF, Rabinowitz JD. 2013. Pyrimidine homeostasis is accomplished by directed overflow metabolism. Nature 500:237–41.

42. Paczia N, Nilgen A, Lehmann T, Gätgens J, Wiechert W, Noack S. 2012. Extensive exometabolome analysis reveals extended overflow metabolism in various microorganisms. Microb Cell Fact 11:122.

43. Pinu FR, Granucci N, Daniell J, Han TL, Carneiro S, Rocha I, Nielsen J, Villas-Boas SG. 2018. Metabolite secretion in microorganisms: the theory of metabolic overflow put to the test. Metabolomics 14:43.

44. Lin H, Hoffmann F, Rozkov A, Enfors SO, Rinas U, Neubauer P. 2004. Change of extracellular cAMP concentration is a sensitive reporter for bacterial fitness in high-cell-density cultures of Escherichia coli. Biotechnol Bioeng 87:602–13.

45. Spari D, Beldi G. 2020. Extracellular ATP as an inter-kingdom signaling molecule: release mechanisms by bacteria and its implication on the host. Int J Mol Sci 21.

46. Harwood CS, Gibson J. 1997. Shedding light on anaerobic benzene ring degradation: a process unique to prokaryotes? J Bacteriol 179:301–9.

47. Schaefer AL, Greenberg EP, Oliver CM, Oda Y, Huang JJ, Bittan-Banin G, Peres CM, Schmidt S, Juhaszova K, Sufrin JR, Harwood CS. 2008. A new class of homoserine lactone quorum-sensing signals. Nature 454:595–9.

48. Gronskiy SV, Zakataeva NP, Vitushkina MV, Ptitsyn LR, Altman IB, Novikova AE, Livshits VA. 2005. The yicM (nepI) gene of Escherichia coli encodes a major facilitator superfamily protein involved in efflux of purine ribonucleosides. FEMS Microbiol Lett 250:39–47.

49. Nygaard P, Saxild HH. 2005. The purine efflux pump PbuE in Bacillus subtilis modulates expression of the PurR and G-box (XptR) regulons by adjusting the purine base pool size. J Bacteriol 187:791–4.

50. McKinlay JB, Oda Y, Rühl M, Posto AL, Sauer U, Harwood CS. 2014. Non-growing Rhodopseudomonas palustris increases the hydrogen gas yield from acetate by shifting from the glyoxylate shunt to the tricarboxylic acid cycle. J Biol Chem 289:1960–1970.

51. D’Souza G, Waschina S, Pande S, Bohl K, Kaleta C, Kost C. 2014. Less is More: Selective advantages can explain the prevalent loss of biosynthetic genes in bacteria. Evolution 68:2559–2570.

