## Supplementary Figures and Tables for "A purine salvage bottleneck leads to bacterial adenine cross-feeding"

**Contents:**

Figures S1-S7
Tables S1-S5
References
Cross-feeding model for testing the validity of adenine diffusion

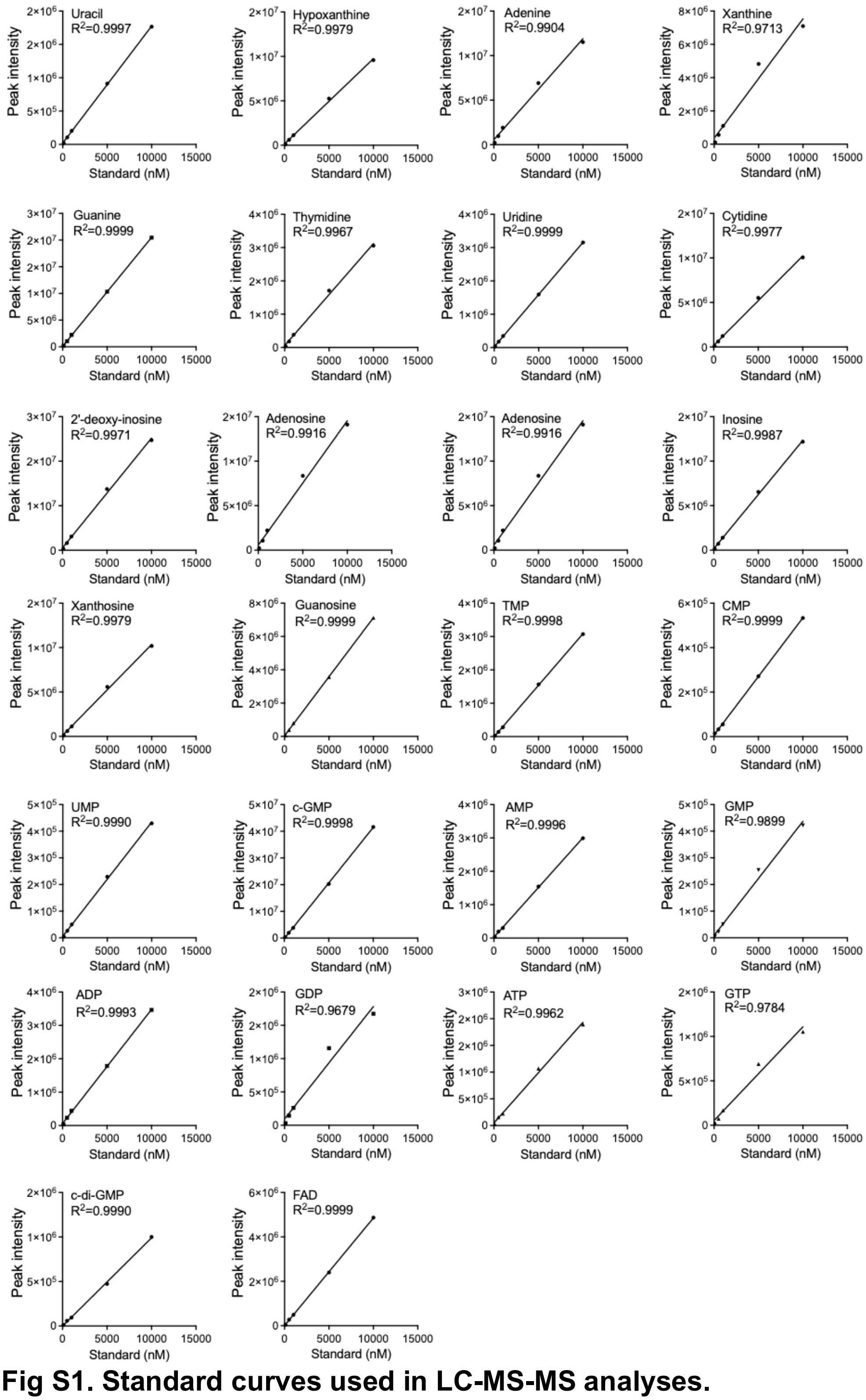

**Fig S1. Standard curves used in LC-MS-MS analyses.**

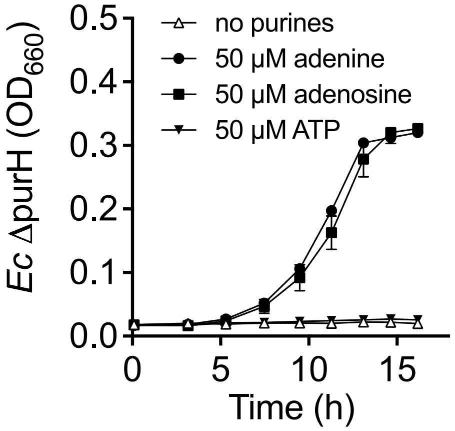

**Fig. S2.** ***E. coli*** **(*Ec*) Δ*purH*, requires purines for growth in monoculture.** Error bars = SD; n=3. Some error bars are smaller than the symbols.

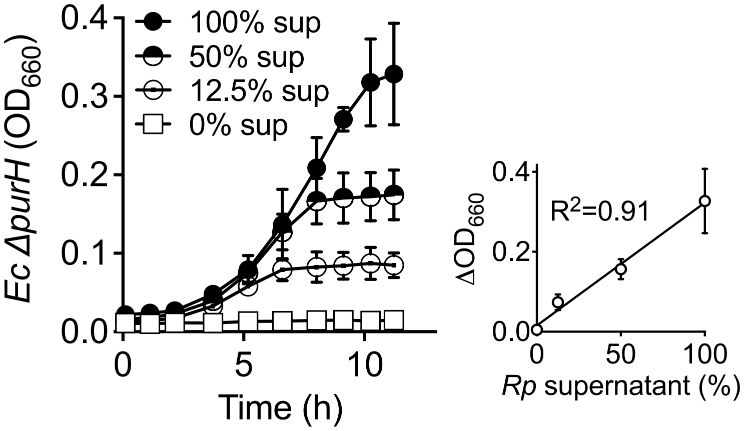

**Fig. S3.** ***R. palustris*** **(*Rp*) CGA4005 supernatants support** ***E. coli*** **(*Ec*) Δ*purH* monoculture growth.** Error bars = SD, n=3. Some error bars are smaller than the symbols. **A.** *E. coli* Δ*purH* monoculture growth curves in media supplemented with difference amounts of *R. palustris* CGA4005 monoculture supernatant **B.** Linear regression of *E. coli* Δ*purH* monoculture growth and the amount of *R. palustris* CGA4005 supernatant provided.

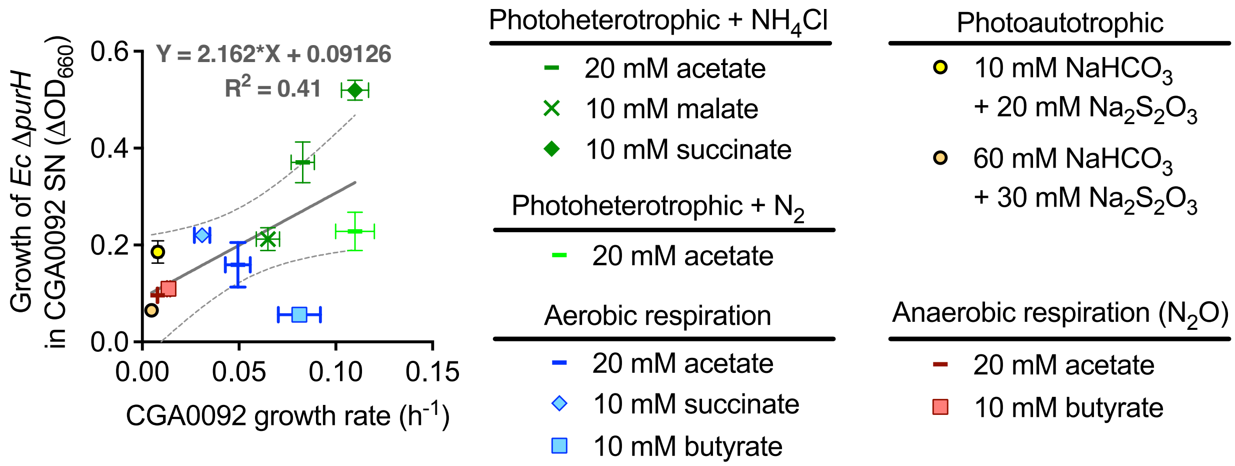

**Fig S4. Purine externalization does not correlate well with CGA0092 growth rate across diverse growth conditions.** Growth of *E. coli* (*Ec*) Δ*purH* in supernatant samples taken from stationary-phase CGA0092 monocultures grown under various growth conditions. Each data point represents the mean of three to six biological replicates +/- SD. Linear regression (gray solid line) +/- 95% confidence intervals (dashed lines) was applied to all samples across all conditions. SN, supernatant.

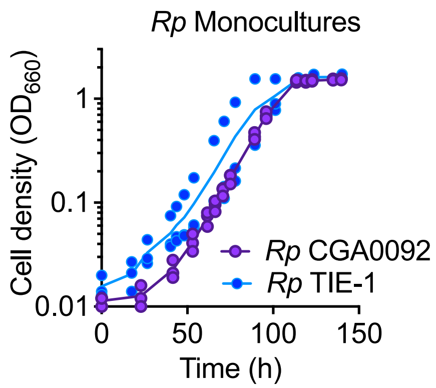

**Fig. S5. Monoculture growth trends are similar for *R. palustris (Rp)* CGA0092 and TIE-1**. Data points from all three biological replicates are shown.

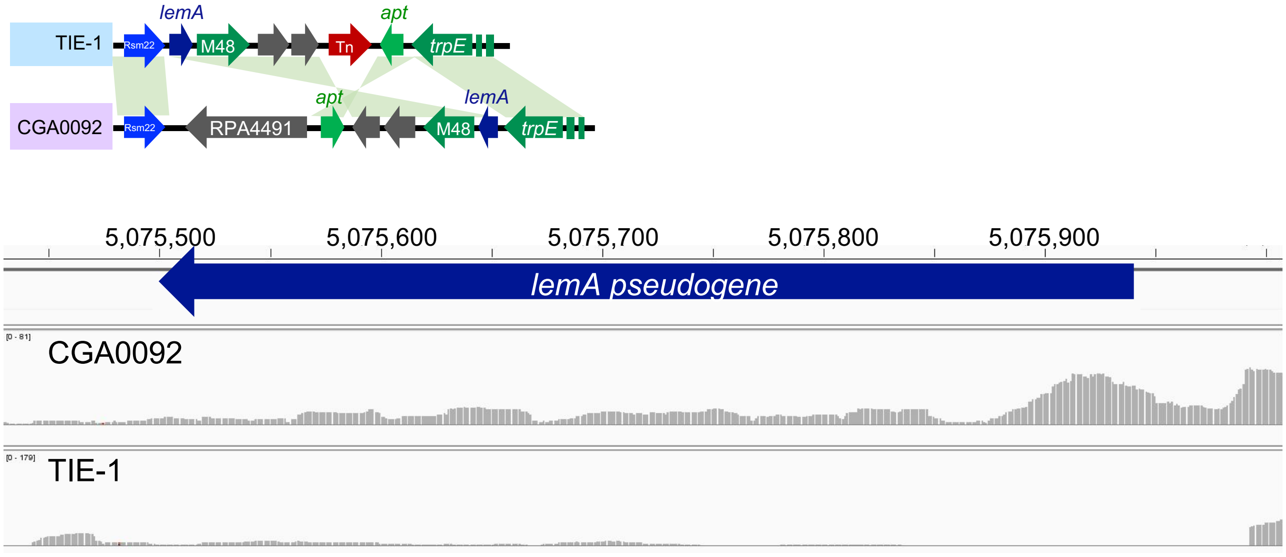

**Fig. S6. CGA0092 exhibits higher *lemA* expression than TIE-1.** Top, orientation of the *lemA*-*apt* cluster in CGA0092 and TIE-1. Bottom, sequencing reads (height of gray bars) across the *lemA* gene in each strain. See the supplementary RNAseq data for the corresponding differential expression values that estimate a 2.2-fold higher transcript levels in CGA0092.

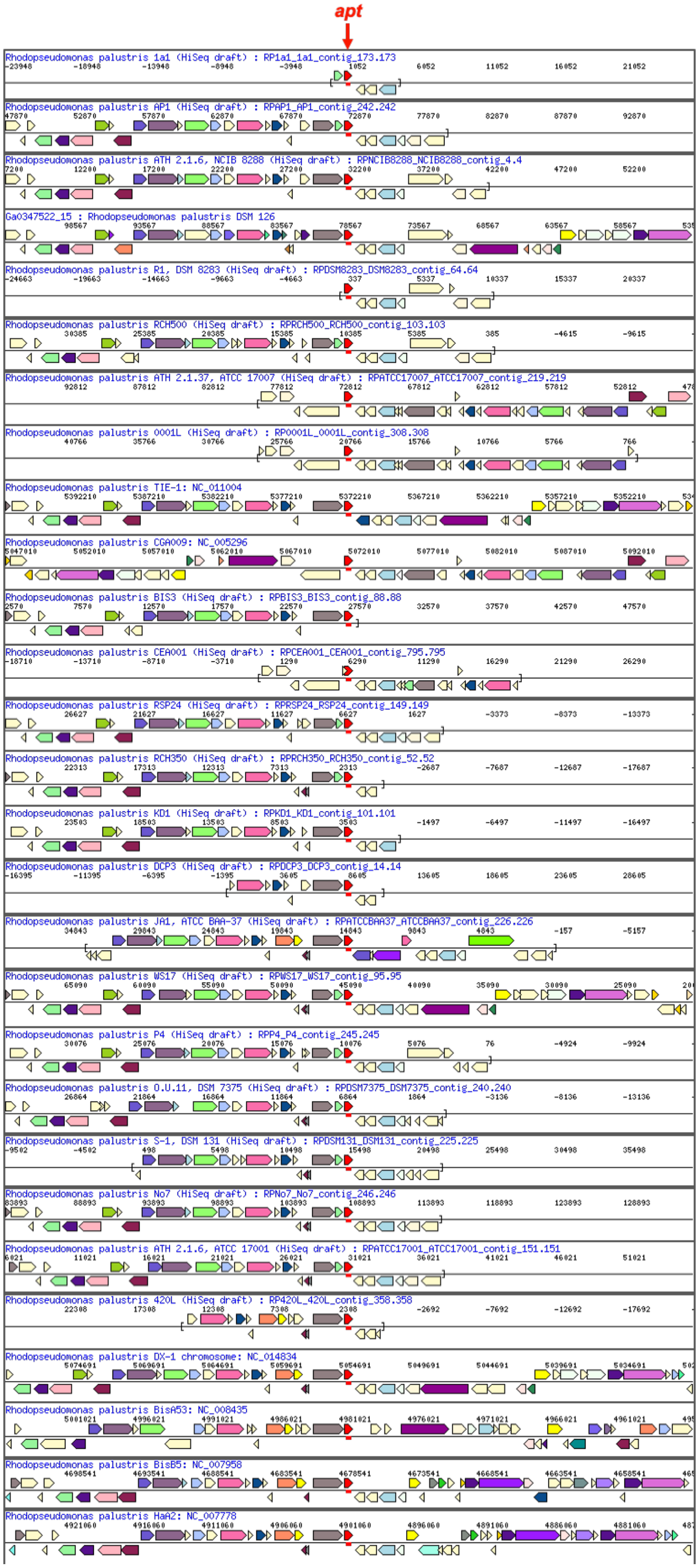

**Fig. S7. Gene neighborhoods for *apt* (red).** Images are from the Joint Genome Institute’s Integrative Microbial Genomes & Microbiomes Gene Neighborhoods tool.

**Table S1. Strains used in this study.**

| Strain | Genotype; 16S rRNA GenBank accession number | Source or Reference |
|---|---|---|
| *R. palustris* | | |
| CGA0092* | Type strain; derivative of CGA001 with spontaneous chloramphenicol resistance | (1) |
| CGA4004 | CGA0092 Δ*hupS* Δ*rpa2750* | (2) |
| CGA4005 | CGA4004 NifA*; a strain with constitutive nitrogenase activity; named Nx in the previous study | (2) |
| TIE-1* | Environmental isolate | (3) |
| DX-1* | Environmental isolate | (4) |
| 1a1* | Environmental isolate | (5) |
| 3ClBen#3 | Environmental isolate; OR656700 | Harwood collection |
| AP1* | Environmental isolate | (5) |
| NCIB8288* | Type strain | DSMZ, (5) |
| DSM126* | Type strain | DSMZ, (5) |
| DSM8283* | Type strain | DSMZ, (5) |
| RCH550 | Environmental isolate; OR656702 | Harwood collection |
| RCH500* | Environmental isolate | (5) |
| ATCC17007* | Type strain | ATCC, (5) |
| 3ClBen#1 | Environmental isolate; OR656699 | Harwood collection |
| 0001L* | Environmental isolate | (5) |
| RCH250 | Environmental isolate; OR656697 | Harwood collection |
| BIS3* | Environmental isolate | (5) |
| RCH300 | Environmental isolate; OR656698 | Harwood collection |
| CEA001* | Environmental isolate | (5) |
| RSP24* | Environmental isolate | (5) |
| RCH450 | Environmental isolate; OR656701 | Harwood collection |
| RCH400 | Environmental isolate; OR659638 | Harwood collection |
| RCH350* | Environmental isolate | (5) |
| KD1* | Environmental isolate | (5) |
| DCP3* | Environmental isolate | (5) |
| RCH150 | Environmental isolate; OR656709 | Harwood collection |
| RCH200 | Environmental isolate; OR656708 | Harwood collection |
| ATCCBAA37* | Type strain | ATCC |
| WS17* | Environmental isolate | (6) |
| P4* | Environmental isolate | (7) |
| Strain#7 | Environmental isolate; OR656710 | Harwood collection |
| DSM7375* | Type strain | DSMZ |
| DSM131* | Type strain | DSMZ |
| No.7* | Environmental isolate | (8) |
| RSP22 | Environmental isolate; OR656707 | Harwood collection |
| RSP21 | Environmental isolate; OR656706 | Harwood collection |
| S55* | Environmental isolate | (8) |
| ATCC17001* | Type strain | ATCC |
| RSP15 | Environmental isolate; OR656705 | Harwood collection |
| RSP10 | Environmental isolate; OR656704 | Harwood collection |
| CGD052 | Environmental isolate; OR656703 | Harwood collection |
| RCH50 | Environmental isolate; OR656696 | Harwood collection |

| | | |
|---|---|---|
| BIS6 | Environmental isolate; OR656711 | (9) |
| 42OL* | Environmental isolate | (10) |
| RSP20 | Environmental isolate; OR656713 | Harwood collection |
| BIS10 | Environmental isolate; OR656712 | (9) |
| ATCC17009 | Type strain; OR656714 | ATCC |
| BisA53* | Environmental isolate | (11) |
| BisB5* | Environmental isolate | (11) |
| HaA2* | Environmental isolate | (11) |
| *E. coli* | | |
| MG1655 | Wild-type K-12 | (12) |
| MG1655 Δ*purH*::Km$^{R}$ | Δ*purH*::Km$^{R}$; mutant allele came from Keio strain JW3970; Referred to as *E. coli* Δ*purH* in the paper. | this study |
| MG1655 Δ*pyrC*::Km$^{R}$ | Δ*pyrC*::Km$^{R}$; mutant allele came from Keio strain JW1049; Referred to as *E. coli* Δ*pyrC* in the paper. | this study |
| NEB10β | DH10B derivative; *Δ(ara-leu) 7697 araD139 fhuA ΔlacX74 galK16 galE15 e14- ϕ80dlacZΔM15 recA1 relA1 endA1 nupG rpsL* (Str$^{R}$) *rph spoT1 Δ(mrr-hsdRMS-mcrBC)* | New England Biolabs |
| JW3970 | Δ*purH*:: *Km$^{R}$*; Keio knockout strain | (13) |
| JW1049 | Δ*pyrC*:: *Km$^{R}$*; Keio knockout strain | (13) |

* 16S rRNA gene sequence is obtained from the Integrated Microbial Genome (IMG) database (https://img.jgi.doe.gov/).

**Table S2. Plasmids used in this study.**

| Plasmids | Genotype or primer sequence (5' to 3'), description | Source or Reference |
|---|---|---|
| pKD46 | Red recombinase expression plasmid; to replace target genes in *E. coli* with PCR products from the Keio collection | (14) |
| pCP20 | Yeast Flp recombinase expression plasmid; to remove the Km resistance cassette in *E. coli* | (15) |
| pBBPgdh | $Gm^R$; pBBR1MCS-5 with a constitutive RPA0944 promoter | (16) |
| pBBPgdh-apt | $Gm^R$; pBBPgdh containing CGA009 *apt* (RPA4492) and the native ribosomal biding site downstream of RPA0944 promoter | This study |

**Table S3. Primers used in this study.**

| Primer | Sequence (5'-3') | Description |
|---|---|---|
| YCC27 | CCC TAT TTG AAC CAG GCA TTA CGC | 5' of Δ*pyrC*::*Km* in JW1049 |
| YCC28 | CGC TGT TTA TCT TCT TTT GTC GCG CC | 3' of Δ*pyrC*::*Km* in JW1049 |
| YCC29 | GCG CAA ACG TTT TCG TTA CAA TGC G | 5' of Δ*purH*::*Km* in JW3970 |
| YCC30 | TGC ATT ACC CGG AGC AAC | 3' of Δ*purH*::*Km* in JW3970 |
| YCC72 | GAG GTC GAC GGT ATC GAT AAG CTC CGT AAC GAA AGA CCC GCG CCG | 5' of RPA4492 (*apt*), overlapping with pBBPgdh |
| YCC73 | AAT TGG AGC TCC ACC GCG GTG GCG GAG AGT TAG GCA GGA GCG AGC | 3' of RPA4492 (*apt*), overlapping with pBBPgdh |
| YCC74 | ATC GGC GCG GGT CTT TCG TTA CGG AGC TTA TCG ATA CCG TCG ACC | Amplify pBBPgdh overlapping with 5' of RPA4492 (*apt*) |
| YCC75 | ATT GAG CTC GCT CCT GCC TAA CTC TCC GCC ACC GCG GTG GAG CTC | Amplify pBBPgdh overlapping with 3' of RPA4492 (*apt*) |
| YCC76 | ATC ATC TTC CGC GAC ATC AC | Forward qPCR primer for *apt* |
| YCC77 | CCT TGT CGA TCT TCG AAC CC | Reverse qPCR primer for *apt* |
| YCC78 | TCG CCG AGG TCG AAG TA | Forward qPCR primer for lemA |
| YCC79 | GTA CAA CAC CGG CAT CCA | Reverse qPCR primer for *lemA* |
| YCC80 | TAA GGG AAC CGT GCA TGT G | Forward qPCR primer for *fixJ* |
| YCC81 | GGA TTC GTA CAG CTT GAC CTC | Reverse qPCR primer for *fixJ* |

**Table S4. Extracellular concentrations of nucleobase-containing compounds in CGA0092 and TIE-1 supernatants.**

| | Concentration (µM / OD) | | | |
|---|---|---|---|---|
| | CGA0092 | | TIE-1 | |
| **Compound** | **Exponential** | **Stationary** | **Exponential** | **Stationary** |
| uracil | 0.58 ± 0.11 | 0.26 ± 0.03 | 0.54 ± 0.06 | 0.37 ± 0.03 |
| hypoxanthine | 0.18 ± 0.02 | 0.12 ± 0.01 | ND | 0.03 ± 0.01 |
| adenine | 17.37 ± 1.67 | 14.36 ± 2.25 | 0.31 ± 0.12 | 0.19 ± 0.03 |
| xanthine | 0.08 ± 0.02 | 0.07 ± 0.04 | 0.07 ± 0.00 | 0.04 ± 0.01 |
| guanine | 0.11 ± 0.03 | 0.07 ± 0.03 | 0.09 ± 0.02 | 0.06 ± 0.01 |
| thymidine | 0.15 ± 0.01 | 0.19 ± 0.05 | 0.22 ± 0.01 | 0.24 ± 0.02 |
| uridine | 0.15 ± 0.05 | 0.11 ± 0.02 | 0.30 ± 0.02 | 0.34 ± 0.05 |
| cytidine | 0.21 ± 0.07 | 0.15 ± 0.02 | 0.37 ± 0.06 | 0.45 ± 0.07 |
| 2'-deoxyinosine | 0.06 ± 0.02 | 0.02 ± 0.00 | 0.04 ± 0.01 | 0.02 ± 0.00 |
| adenosine | 0.52 ± 0.20 | 0.49 ± 0.08 | 0.63 ± 0.13 | 0.56 ± 0.05 |
| 2'-deoxyguanosine | 0.07 ± 0.02 | 0.06 ± 0.02 | 0.06 ± 0.01 | 0.04 ± 0.00 |
| inosine | 0.13 ± 0.02 | 0.12 ± 0.04 | 0.07 ± 0.01 | 0.05 ± 0.00 |
| xanthosine | 0.12 ± 0.02 | 0.10 ± 0.01 | 0.13 ± 0.01 | 0.11 ± 0.01 |
| guanosine | 0.17 ± 0.06 | 0.12 ± 0.02 | 0.17 ± 0.03 | 0.15 ± 0.01 |
| TMP | 0.09 ± 0.03 | 0.05 ± 0.01 | 0.08 ± 0.02 | 0.07 ± 0.01 |
| CMP | 0.17 ± 0.06 | 0.10 ± 0.01 | 0.13 ± 0.02 | 0.23 ± 0.01 |
| UMP | 0.19 ± 0.03 | 0.11 ± 0.01 | 0.21 ± 0.03 | 0.33 ± 0.02 |
| c-GMP | 0.07 ± 0.02 | 0.03 ± 0.01 | 0.06 ± 0.01 | 0.04 ± 0.00 |
| AMP | 0.10 ± 0.03 | 0.08 ± 0.02 | 0.13 ± 0.02 | 0.20 ± 0.01 |
| GMP | 0.08 ± | 0.06 ± 0.00 | 0.16 ± 0.03 | 0.20 ± 0.04 |
| ADP | 0.21 ± 0.06 | 0.11 ± 0.02 | 0.21 ± 0.04 | 0.19 ± 0.02 |
| GDP | 0.12 ± | 0.07 ± 0.02 | 0.13 ± 0.04 | 0.13 ± 0.04 |
| ATP | 0.35 ± 0.21 | 0.20 ± 0.03 | 0.43 ± 0.07 | 0.05 ± 0.02 |
| GTP | ND | ND | 0.09 ± | 0.05 ± 0.01 |
| c-di-GMP | 0.08 ± 0.03 | 0.03 ± 0.01 | 0.07 ± 0.02 | 0.04 ± 0.01 |
| FAD | 0.10 ± 0.03 | 0.04 ± 0.00 | 0.08 ± 0.01 | 0.04 ± 0.01 |

ND, not detected. Values are means ± SD; n= 1-3. Triplicate samples were run in each case but not all compounds were detected in each sample. Values without SD indicate that a compound was only detected in one of the three replicates.

**Table S5. Intracellular concentrations of nucleobase-containing compounds in CGA0092 and TIE-1 supernatants.**

| | Concentration (µM) | | | |
|---|---|---|---|---|
| | CGA0092 | | TIE-1 | |
| Compound | Exponential | Stationary | Exponential | Stationary |
| uracil | 30.0 ± 8.4 | 19.5 ± 4.9 | 31.2 ± 7.1 | 44.2 ± 2.2 |
| hypoxanthine | 29.5 ± 7.7 | 24.1 ± 5.7 | ND | ND |
| adenine | 1516.8 ± 645.0 | 1467.5 ± 554.3 | 17.0 ± 3.1 | 13.7 ± 4.5 |
| xanthine | ND | 12.6 ± 4.2 | 11.8 ± | 7.6 ± |
| guanine | 22.6 ± 3.4 | 20.5 ± 3.6 | 22.9 ± 6.3 | 21.7 ± 1.0 |
| thymidine | 23.3 ± 8.0 | 26.8 ± 3.6 | 27.8 ± 11.0 | 42.2 ± 3.1 |
| uridine | 18.4 ± 1.8 | 14.7 ± 1.9 | 28.8 ± 7.4 | 46.8 ± 5.1 |
| cytidine | 35.9 ± 4.0 | 32.9 ± 6.8 | 53.2 ± 14.5 | 102.0 ± 11.8 |
| 2'-deoxyinosine | 10.8 ± 2.4 | 5.1 ± 0.6 | 9.6 ± 1.8 | 4.7 ± 0.1 |
| adenosine | 67.1 ± 11.7 | 67.8 ± 1.9 | 59.9 ± 25.6 | 105.9 ± 16.3 |
| 2'-deoxyguanosine | 15.9 ± 3.0 | 14.2 ± 4.2 | 15.1 ± 3.1 | 11.4 ± 1.0 |
| inosine | 24.6 ± 6.1 | 22.4 ± 3.1 | 16.6 ± 3.7 | 13.2 ± 0.3 |
| xanthosine | 24.7 ± | 20.3 ± 2.2 | 22.6 ± 5.6 | 26.4 ± 3.3 |
| guanosine | 32.9 ± 4.4 | 31.3 ± 5.1 | 32.0 ± 9.8 | 47.5 ± 2.3 |
| TMP | 25.0 ± 8.5 | 13.8 ± 0.7 | 21.5 ± 6.9 | 17.5 ± 2.2 |
| CMP | 54.0 ± 22.1 | 27.0 ± 5.0 | 36.4 ± 5.7 | 84.1 ± 22.2 |
| UMP | 47.4 ± 13.0 | 41.0 ± 7.3 | 53.2 ± 10.1 | 97.3 ± 7.4 |
| c-GMP | 17.8 ± 4.0 | 10.0 ± 0.5 | 17.5 ± 3.7 | 10.6 ± 0.3 |
| AMP | 84.3 ± 20.3 | 142.9 ± 22.1 | 105.7 ± 24.7 | 167.6 ± 9.5 |
| GMP | 40.5 ± | 33.7 ± 6.1 | 57.1 ± 18.6 | 58.2 ± 9.1 |
| ADP | 168.6 ± 12.6 | 244.5 ± 69.9 | 177.0 ± 3.7 | 217.9 ± 38.1 |
| GDP | 65.2 ± | 54.0 ± 8.6 | 63.3 ± 8.2 | 82.0 ± 8.9 |
| ATP | 356.8 ± 97.0 | 406.9 ± 187.9 | 301.9 ± 49.3 | 319.1 ± 46.7 |
| GTP | 59.0 ± 11.8 | 44.6 ± 8.4 | 53.4 ± 5.7 | 81.2 ± 19.0 |
| c-di-GMP | 21.8 ± 1.9 | 13.5 ± 3.3 | 18.1 ± 6.5 | 12.8 ± 1.3 |
| FAD | 38.4 ± 10.7 | 29.1 ± 8.4 | 33.6 ± 3.7 | 32.5 ± 9.0 |

ND, not detected. Values are means ± SD; n= 1-3. Triplicate samples were run in each case but not all compounds were detected in each sample. Values without SD indicate that a compound was only detected in one of the three replicates.

#### Supplementary Figure and Table References:

1. Mazny BE, Sheff OF, LaSarre B, McKinlay A, McKinlay JB. 2023. Complete genome sequence of *Rhodopseudomonas palustris* CGA0092 and corrections to the *R. palustris* CGA009 genome sequence. Microbiol Resour Announc 12:e0128522.
2. LaSarre B, McCully AL, Lennon JT, McKinlay JB. 2017. Microbial mutualism dynamics governed by dose-dependent toxicity of cross-fed nutrients. ISME J 11:337-348.
3. Jiao Y, Kappler A, Croal LR, Newman DK. 2005. Isolation and characterization of a genetically tractable photoautotrophic Fe(II)-oxidizing bacterium, *Rhodopseudomonas palustris* strain TIE-1. Appl Environ Microbiol 71:4487-96.
4. Xing D, Zuo Y, Cheng S, Regan JM, Logan BE. 2008. Electricity generation by *Rhodopseudomonas palustris* DX-1. Environ Sci Technol 42:4146-51.
5. Fixen KR, Oda Y, Harwood CS. 2016. Clades of photosynthetic bacteria belonging to the genus *Rhodopseudomonas* show marked diversity in light-harvesting antenna complex gene composition and expression. mSystems 1:e00006-15.
6. Kamal VS, Wyndham RC. 1990. Anaerobic phototrophic metabolism of 3-chlorobenzoate by *Rhodopseudomonas palustris* WS17. Appl Environ Microbiol 56:3871-3.
7. Jung G, Jung H, Kim J, Ahn Y, Park S. 1999. Isolation and characterization of *Rhodopseudomonas palustris* P4 which utilizes CO with the production of $H_2$. Biotechnol Lett 21:525-529.
8. Inui M, Roh JH, Zahn K, Yukawa H. 2000. Sequence analysis of the cryptic plasmid pMG101 from *Rhodopseudomonas palustris* and construction of stable cloning vectors. Appl Environ Microbiol 66:54-63.
9. Oda Y, Meijer WG, Gibson JL, Gottschal JC, Forney LJ. 2004. Analysis of diversity among 3-chlorobenzoate-degrading strains of *Rhodopseudomonas palustris*. Microb Ecol 47:68-79.
10. Carlozzi P, Pushparaj B, Degl'Innocenti A, Capperucci A. 2006. Growth characteristics of *Rhodopseudomonas palustris* cultured outdoors, in an underwater tubular photobioreactor, and investigation on photosynthetic efficiency. Appl Microbiol Biotechnol 73:789-95.
11. Oda Y, Larimer FW, Chain PSG, Malfatti S, Shin MV, Vergez LM, Hauser L, Land ML, Braatsch S, Beatty JT, Pelletier DA, Schaefer AL, Harwood CS. 2008. Multiple genome sequences reveal adaptations of a phototrophic bacterium to sediment microenvironments. Proc Natl Acad Sci USA 105:18543-18548.
12. Blattner FR, Plunkett G, 3rd, Bloch CA, Perna NT, Burland V, Riley M, Collado-Vides J, Glasner JD, Rode CK, Mayhew GF, Gregor J, Davis NW, Kirkpatrick HA, Goeden MA, Rose DJ, Mau B, Shao Y. 1997. The complete genome sequence of *Escherichia coli* K-12. Science 277:1453-62.
13. Baba T, Ara T, Hasegawa M, Takai Y, Okumura Y, Baba M, Datsenko KA, Tomita M, Wanner BL, Mori H. 2006. Construction of *Escherichia coli* K-12 in-frame, single-gene knockout mutants: the Keio collection. Mol Syst Biol 2:2006.0008.
14. Datsenko KA, Wanner BL. 2000. One-step inactivation of chromosomal genes in *Escherichia coli* K-12 using PCR products. Proc Natl Acad Sci USA 97:6640-6645.

15. Cherepanov PP, Wackernagel W. 1995. Gene disruption in *Escherichia coli*: TcR and KmR cassettes with the option of Flp-catalyzed excision of the antibiotic-resistance determinant. Gene 158:9-14.
16. McKinlay JB, Harwood CS. 2010. Carbon dioxide fixation as a central redox cofactor recycling mechanism in bacteria. Proc Natl Acad Sci USA 107:11669-11675.

**Cross-feeding model for testing the validity of adenine diffusion**

````
---
title: "R. palustris- E. coli interaction Model with purine transfer"
authors: "James B. McKinlay"
date: "`r format(Sys.time(), '%d %B, %Y')`"
header-includes:
  - \usepackage{array}
output: pdf_document
geometry: margin=2.54cm
---

## Overview
Monod model simulating population and extracellular metabolism in a coculture of an
E.coli purine auxotroph and R. palustris. Under anaerobic conditions, E. coli ferments
glucose to excreted products, some of which serve as a carbon source for R. palustris.
R. palustris excretes adenine which is required by the E. coli auxotroph. The following
attempts to captures these basic features using a set of coupled differential equations.
To simplify the output, the model does not include fermentation products that do not
contribute to acidification or cross-feeding, namely ethanol, H2, and CO2.

## Set working directory
```{r}
rm(list=ls()) # clears your working directory
getwd() # tells you your current working directory
setwd("<enter working directory from getwd here>") # sets your working directory
```

## Install packages
```{r}
#install.packages("deSolve") # run this the first time by removing the '#'
require("deSolve") # installs any dependencies
```

## Mutualism function
```{r}
mutfun <- function(times, y, parms) {
  with(as.list(c(y, parms)), {

    uEc <- uEcMax*(G/(Kg+G))*(P/(Kp+P))*(bEc/(bEc+10^(f+C)))

     # E.coli growth rate (h^-1); controlled by Monod function where G = glucose
concentration, Kg = half saturation constant for G, P = purine (adenine) concentration,
Kp = half saturation constant for P; bEc/(bEc+10^(f+C)) is a sigmoidal function
resembling a buffer titration curve where accumulation of formate (f) and consumable
organic acids (C) will decrease the growth rate
````
````

```
  uRp <- uRpMax*(C/(Kc+C))*(bRp/(bRp+10^(f+C)))

  # R. palustris growth rate (h^-1); controlled by Monod function where C = consumable organic acid (OAcs) concentration, Kc = half saturation constant for C; growth rate is also controlled by acid accumulation as described for the E. coli growth rate

 dGdt <- -uEc*Ec/Yg   - Ec*(G/(Kg+G))*(10/(10+1.09^(1000*uEc)))*(bEc/(bEc+10^(f+C)))*((100/(100+6^C))*(rc+ rf))

# change in glucose (mM/h); the 2nd function describes growth independent metabolism as described by McCully et al, 2017, Environ Microbiol; applying the Monod and sigmoidal acid accumulation functions constrains growth independent metabolism to be dependent on glucose availability and permissive acid levels

 dPdt <-  -uEc*Ec/Yp + Rp*IP*Perm*SA*(C/(Kc+C))*(bRp/(bRp+10^(f+C)))

  # change in purine (adenine) conc (mM/h); Ec = E. coli cells/ml; Yp = growth yield on adenine; Rp = Rpalustris cells/ml; IP = intracellular adenine conc (umol/cm^3); Perm = adenine permeability coefficient (cm/s); SA = cellular surface area (cm^2/cell) ; applying the Monod and sigmoidal acid accumulation functions constrains diffusion to be dependent on organic acid availability and permissive acid levels

  dCdt <- -uRp*Rp/Yc  + uEc*Ec*Fc + Ec*(G/(Kg+G))*(10/(10+1.09^(1000*uEc)))*(bEc/(bEc+10^(f+C)))*(rc*(100/(100+6^C)))# +rc_mono)

# change in organic acids (OAcs; mM/h); ; Yc = Rp growth yield on OAcs; Fc = OAcs excretion level (umol/Ec cell); the 2nd function describes growth independent metabolism as described by McCully et al, 2017, Environ Microbiol;

  dFdt <- uEc*Ec*Ff + Ec*(G/(Kg+G))*(10/(10+1.09^(1000*uEc)))*(bEc/(bEc+10^(f+C)))*(rf*(100/(100+6^C)))# + rf_mono)

# change in formate (mM/h); Ff = formate excretion (umol/Ec cell); the 2nd function describes growth independent metabolism as described by McCully et al, 2017, Environ Microbiol;

  dRpdt <-Rp*uRp

#change in Rp cell density (cells/ml)

  dEcdt <- Ec*uEc
```

~~~~
 #change in Ec cell density (cells/ml)

    return(list(c(dGdt, dPdt, dCdt, dFdt, dRpdt, dEcdt)))
  })
}
```

## Parameters and initial conditions
```{r}

### Parameters
parms <- c(

###### E.coli ####

uEcMax = 0.28, # E. coli max growth rate (h^-1; McCully et al, 2017, Environ Microbiol)

Yg = 4.4*10^6,# E. coli cells/umol; based on time course data in Fig 2

Yp = 6*10^9, # E. coli cells/umol adenine; based on bioassay std curve in Fig 4

Fc = 3.44*10^-7, # umol organic acids / E. coli cell; Based on Fig 2: the sum of organic
acids left over in coculture (2.20*10^-7 umol/ E.coli cell) + what was estimated to be
assimilated by R. palustris (1.24*10^-7)

Ff = 2*10^-7, # umol/cell; based on what is left over in coculture

Kg = 0.02, # E. coli half saturation constant for glucose (mM; Buhr et al 1992 J Biol
Chem)

Kp = 0.004, # E. coli half saturation constant for adenine (mM; 0.004 = average for PurP
(0.001) and YicO (0.0065); Papakostas et al. 2013. JBC 288: P36827-36840

rc = 300*10^-11, # mM glucose converted to organic acids per E. coli cell/ml per h in the
absence of growth (umol/cell/h; McCully et al, 2017, Environ Microbiol)

rf = 50*10^-11,# mM glucose converted to formate per E. coli cell/ml/h in the absence of
growth (umol/cell/h; rounded up from McCully et al, 2017, Environ Microbiol)

###### R. palustris #####
uRpMax = 0.0875, #R. palustris max growth rate with NH4Cl (h^-1 )

Yc = 2*10^8, # R. palustris growth yield on organic acids (cells/umol OAcs consumed;
based on values from this study)

Kc = 0.01, # half saturation constant for organic acids (mM; assumed)
~~~~

````

Perm = 1.38e-5, # adenine permeability coefficient (cm/s; Xiang and Anderson, 1994, J Membrane Biol, 140, 111-122)

SA = 5.1e-8, # R. palustris surface area (cm^2; based on a 3.25 um cell length; LaSarre 2018 mBio, Fig 3F) Smaller and larger SA values were upper and lower bounds as follows: 4.24e-8 cm^2 for 2.7 um cell length; 5.97e-8 cm^2 for 3.8 um cell length

IP = 1.5, # Intracellular adenine concentration (mM or umol/cm3; 1.5 mM for CGA009; 0.02 mM for TIE-1)

bEc = 1*10^43, # resistance of E. coli to acid; reflects medium buffering (McCully et al, 2017, Environ Microbiol)

bRp = 1*10^32) # resistance of R. palustris to acid; reflects medium buffering (McCully et al, 2017, Environ Microbiol)

# initial concentrations:
init <- c(
	G = 22.5, # Glucose (mM)
	P = 0.00000001, # adenine (mM); non-zero to initiate growth
	C = 0.00000001, # OAcs (mM); non-zero to initiate growth; set to 10 for Rp monocultures
	f = 0, # Formate (mM)
	Rp = 3000000, (cells/ml; based on Fig 2B)
	Ec = 6000000) (cells/ml; based on Fig 2B); set to 0 for Rp monocultures

# Time range and time step
Tmax = 200 # hours
TimeStep = 1 # hours
Time = seq(0, Tmax, by = TimeStep)
```

#### Output
```{r}
out <-  ode(y = init, times = Time, func = mutfun, parms = parms, method = "lsode")
#head(out) # prints first 6 time points
#(out) #prints all time points
tail(out) # prints last 6 time points
write.csv(out, file = "out.csv") # creates comma-delimited file that can be pasted into spreadsheet for graphing
```

#### Plotting
```{r}

````

````
plot(out[,1], out[,7], xlab = "time (hr)", ylab = "Ec", type = "l", las = 1,
    col = "orange", lwd = 4, ylim = c(0, 1.1*max(out[,7]))) # plot P2

par(new=T)
plot(out[,1], out[,6], xlab="", ylab ="", type = "l",las = 1,
    col = "purple", ylim=c(0,1*max(out[,6])) )
axis(4, ylim=c(0,1.2*max(out[,6])), lwd=2, line=3.5)
mtext(4, text="Rpal", line=5.5) # plot R. palustris biomass

par(new=T)
plot(out[,1], out[,4], xlab="", ylab ="",  type = "l",las = 1,
    col = "blue", ylim=c(0,1.3*max(out[,4])) )
axis(4, ylim=c(0,1.2*max(out[,4])), lwd=2, line=3.5)
mtext(4, text="Oacs", line=5.5) # plot organic acids

par(new=T)
plot(out[,1], out[,5], xlab="", ylab ="",  type = "l",las = 1,
   col = "green", ylim=c(0,1.3*max(out[,5])) )
axis(4, ylim=c(0,0.1*max(out[,5])), lwd=2, line=3.5)
mtext(4, text="Formate", line=5.5) # plot formate

par(new=T)
plot(out[,1], out[,3], xlab="", ylab ="",  type = "l",las = 1,
    col = "red", ylim=c(0,1*max(out[,3])) )
axis(4, ylim=c(0,1*max(out[,3])), lwd=4, line=3.5)
mtext(4, text="purine", line=5.5) # plot purine
par(new=T)

plot(out[,1], out[,2], xlab="", ylab ="",  type = "l",las = 1,
    col = "black", ylim=c(0,1*max(out[,2])) )
axis(4, ylim=c(0,1*max(out[,2])), lwd=3, line=3.5)
mtext(4, text="glucose", line=5.5) # plot glc
```
````
````
